# Quantitative prediction of nonsense-mediated mRNA decay across human genes by genomic language model and large-scale mutational scanning

**DOI:** 10.64898/2026.03.24.714003

**Authors:** Marcell Veiner, Ignasi Toledano, Guillermo Palou-Márquez, Ben Lehner, Fran Supek

## Abstract

The molecular consequences of protein truncating variants depend strongly on whether their transcripts are eliminated by nonsense-mediated mRNA decay (NMD), yet NMD is still predicted largely from a small set of binary positional rules. How individual premature termination codons (PTCs) engage NMD across genes and transcript contexts therefore remains incompletely resolved. Here we integrated endogenous allele-specific PTC expression from large-scale genomic data, mRNA language-model prediction and high-throughput mutational scanning to revisit the rules that govern mammalian NMD. Using allele-specific expression measurements from large human cohorts, we trained NMDetective-AI on ∼14,000 somatic PTCs, and after testing it on ∼1,800 germline PTCs, found that it improves on previous models, with its accuracy approaching the reproducibility of the underlying measurements. We then generated experimental maps of NMD with deep mutational scanning, including ∼450 PTCs nearby the 50-nt penultimate exon boundary, ∼950 engineered PTCs across 9 exon lengths to resolve long-exon escape, and ∼11k PTCs across 139 genes to quantify and refine start-proximal evasion. Our results support the positional logic of mammalian NMD yet show that this logic is implemented quantitatively: the classical rules resolve into graded, gene-dependent response curves whose boundaries are shaped by transcript architecture and modulated by local sequence context. Applying this framework to population, disease and cancer datasets further identifies genes in which NMD is predicted to aggravate or ameliorate the effects of truncating variants, providing a basis for variant interpretation and for prioritizing NMD-directed therapies.

## Introduction

Nonsense-mediated mRNA decay (NMD) is a translation-dependent quality control pathway that actively degrades transcripts harboring premature termination codons (PTCs) due to mutation or altered splicing, thereby preventing the accumulation of truncated, potentially harmful peptides. Beyond surveillance of pathological transcripts, NMD also controls a pervasive regulatory network, constitutively degrading ∼5-20% of the physiological transcriptome in homeostatic conditions^1–4^.

The biophysical decision “algorithm” that dictates whether a specific PTC-containing transcript is degraded by NMD or evades detection (allowing translation of a truncated protein) relies strongly on the longitudinal arrangement of features along the transcript, specifically the position of the terminating ribosome relative to downstream exon junctions. In mammalian cells, the splicing machinery deposits an exon junction complex (EJC) ∼20–24 nucleotides upstream of an exon-exon junction. According to the classical “50-nucleotide rule,” if a ribosome stalls at a PTC located more than 50–55 nucleotides upstream of the final exon-exon junction, the downstream EJC is not displaced by the terminating ribosome. This retained EJC subsequently recruits UPF and SMG surveillance factors (e.g. UPF1, SMG1, SMG6, SMG7), triggering endonucleolytic and exonucleolytic decay of the mRNA. Conversely, PTCs located in the final exon, or within the distal ∼50 nucleotides of the penultimate exon, typically fail to trigger the NMD machinery^5–9^ because the terminating ribosome clears all downstream EJCs. Additional exceptions to canonical EJC-dependent decay have also been identified, including start-proximal NMD evasion in the first ∼100-200 nts of the coding sequence, often driven by translation reinitiation at downstream AUG codons^6,10–13^ and EJC-independent NMD triggered by long 3′ untranslated regions (UTRs)^14–16^.

Historically, these topological NMD rules were painstakingly elucidated through low-throughput, locus-specific functional assays on individual disease genes such as *HBB* (e.g.^17^ *BRCA1* (e.g.^18^), or *DMD* (e.g.^19^). The advent of massive, pan-cancer and population-scale genomic datasets (e.g., TCGA, GTEx, gnomAD) allowed researchers to evaluate NMD rules genome-wide by associating gene expression differences with presence of a PTC^6,9,20,21^. These large-scale analyses confirmed that while the canonical NMD rules (such as the 50-nt boundary) are statistically robust at a global scale, they are lacking as quantitative predictors, leaving a substantial fraction – approximately one-third – of the variance in NMD efficiency across PTCs unexplained^22^. The above highlights the limitations of treating NMD as a simple binary switch; instead, the likely case is that NMD efficiency is a continuous quantity, with NMD rules having “blurry” boundaries, and additionally modulated by local sequence context and other complex biophysical features that have historically been abstracted away.

In addition to the canonical 50-nucleotide boundary, several other NMD rules have been proposed largely on the basis of bioinformatic inference. For instance, the “long-exon rule”, which posits that PTCs residing in exons >400 nt partially escape NMD speculatively due to the spatial dissociation of the terminating ribosome from downstream EJCs – was derived from cancer genomic dataset analysis^6^ and independently replicated in other genomic datasets^20,23^. However, this lacks systematic experimental support, leaving unclear possible confounding effects and also the exact contributions of PTC distances to 5’ and 3’ exon ends towards determining NMD activity. Similarly, the “start-proximal rule”, reported from single-gene studies and recapitulated in genomics observational data analyses^6,20,21,23,24^, still has limited scope of experimental validation: the shape of the decrease in NMD activity towards the 5’ coding region is not well characterized, as well as the quantitative relevance of translation reinitiation as an NMD escape mechanism.

As a conceptual limitation, it remains unknown to what extent these NMD rules vary across the transcriptome. A priori, it does not seem likely that the boundaries of local NMD-evading regions would be universally fixed at a precise nucleotide distance; rather, they may “wobble” from gene to gene, for instance due to RNA structure, cis-regulatory elements or ribosome dynamics, and the extent of this between-gene variability is unknown. To model this presumably sequence-dependent variability, deep learning architectures – specifically DNA/RNA genomic language models (reviewed in^25^) – offer a powerful paradigm. By extracting the hidden grammar of the transcriptome from vast evolutionary datasets^26–28^ these models can in principle identify complex sequence motifs and interactions; one example thereof would be positioning relative to exon-exon junctions, which is critical for NMD modelling. Ultimately, however, even sophisticated *in silico* predictions based on transcript sequence remain correlative, unless supported by (ideally high-throughput) experimental validation that directly observes effects of various introduced PTCs on NMD activity, in context of specific placement.

Achieving high-resolution, accurate models of NMD efficiency applicable to every transcript is a necessity for translational applications in genetic medicine. Accurate NMD predictions are needed to resolve the molecular impact of putatively pathogenic nonsense and frameshift variants^22,23,29–32^ and to clarify the dual, often paradoxical role of NMD in genetic disease variants, including cancer driver mutations (reviewed in^33–35^). For some loci, NMD clears transcripts that would otherwise encode harmful dominant-negative or neomorphic proteins; in others, NMD degrades transcripts that could have yielded partially-functional protein truncations. Disentangling the former case of NMD-ameliorated genetic disease alleles, from the latter case of NMD-aggravated PTC alleles is a prerequisite for inferring whether a patient will safely benefit from NMD inhibitors and/or PTC readthrough therapies^36–41^.

Here, we integrate pan-tissue genomic and transcriptomic datasets to map the quantitative landscape of NMD efficiency at base-pair resolution. By leveraging allele-specific expression as a precise readout of transcript levels contingent on a PTC, we train NMDetective-AI, a deep learning framework built upon a foundational RNA language model. To systematically validate the model’s predictions and empirically dissect the physical boundaries of local NMD evasion, including the long-exon and start-proximal rules, we couple our computational inferences with a comprehensive high-throughput deep mutational scanning (DMS) interrogating >100 genes in human cells. Finally, we apply this integrated framework to clinical and cancer genomic databases to clarify the pathogenicity of germline and somatic nonsense variants. By stratifying disease genes into NMD-aggravated and NMD-ameliorated categories, we define a predictive genomic map that clarifies potential for benefit from pharmacological NMD modulation guided by the genomic sequence.

## Results

### Quantifying NMD efficiency of nonsense variants in somatic human tissues at a large scale

A comprehensive, accurate training dataset is essential for developing a highly performant machine learning model, and we aimed to assess premature termination codons (PTCs) expressed across a variety of human tissues for their visibility to NMD silencing at the mRNA level. By integrating whole-exome sequencing (WES) variant calls and RNA-seq data, we assembled a large-scale dataset of tumor tissues, in which somatic mutations and germline variants causing PTCs were considered, and of healthy human tissues, in which germline PTCs were analyzed for their association with mRNA levels (Fig. 1). In particular, we assessed NMD efficiency for each PTC observed in TCGA (9,752 individuals, 8,786 tumor samples, 31 cancer types) and GTEx^42^ (838 individuals, 15,247 healthy samples, 56 tissues) using allele-specific expression (ASE), as the negative log2 of the ratio in mRNA expression of the mutant allele relative to the wild-type allele (Methods).

**Figure 1:**
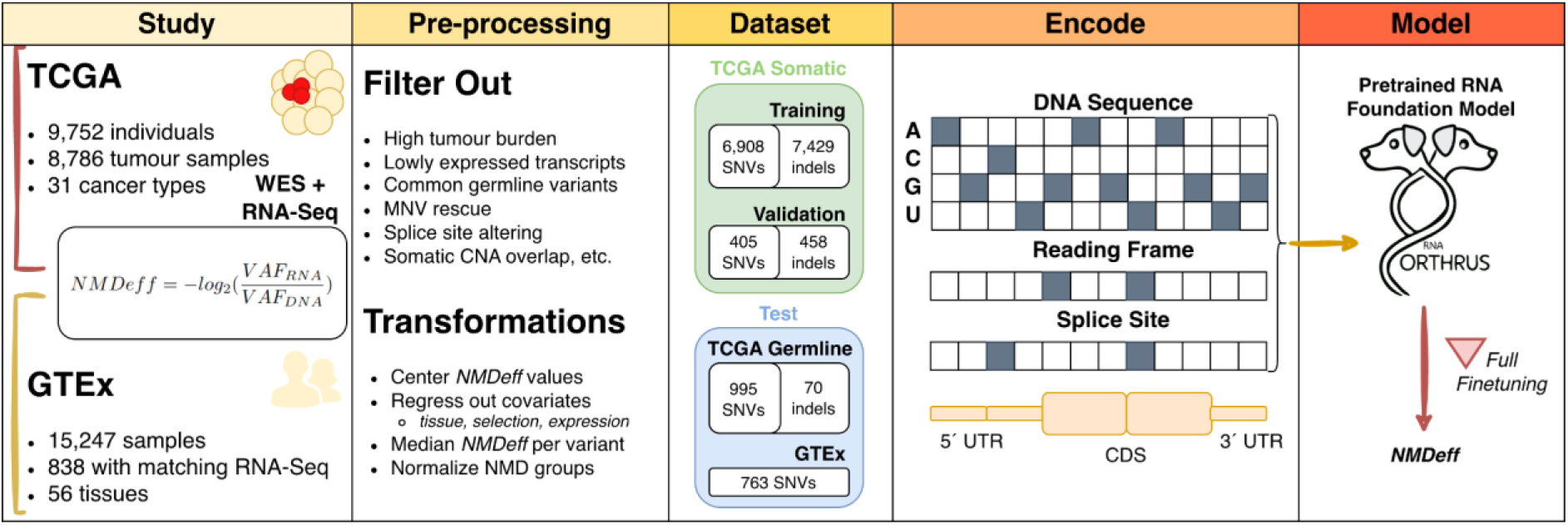
Overview of the NMD efficiency datasets and NMDetective-AI. Nonsense-mediated decay (NMD) efficiency (NMDeff) was quantified for premature termination codons (PTCs) identified in TCGA (somatic and germline) and GTEx (germline) cohorts. NMDeff is calculated as the negative log2 of the ratio in mRNA expression of the mutant allele relative to the wild-type allele. Pre-processing: Raw PTC calls were filtered to remove confounders such as high tumor mutational burden, somatic copy number alterations, and multi-nucleotide variants (see Methods). For modelling, values were further transformed by regressing out technical covariates and normalizing against canonical NMD-triggering and evading controls. Dataset: The curated dataset was partitioned into training/validation sets by chromosomes (TCGA somatic) and independent test sets (TCGA germline and GTEx), comprising both single-nucleotide variants (SNVs) and indels. Encode: Transcript sequences (5’ UTR, CDS, and 3’ UTR) were represented as 6-track one-hot encodings, incorporating the DNA sequence, the reading frame/CDS boundaries, and splice site positions. Model: The NMDetective-AI model, built upon the pretrained Orthrus foundation model, underwent full finetuning to predict normalized NMDeff from the multi-track sequence input.

We applied standard stringent quality filters to the PTC calls (Fig. S1), such as filtering out variants in regions affected by somatic copy number alterations, lowly expressed genes, common germline variants, multi-nucleotide variants (MNV) that may be rescued by adjacent substitutions and ambiguous NMD classification between transcripts of the same gene^1,20,32^, and others (see Methods for exhaustive list of filtering steps; summary in Fig. 1; Fig S1). The resulting dataset comprises 22,535 somatic TCGA PTCs (9,059 nonsense SNVs and 13,476 frameshifting indels), 1,550 germline TCGA PTCs (1,401 nonsense and 149 indels), and 11,335 germline GTEx PTCs (all nonsense).

As a check on the dataset quality, we considered PTCs classified as NMD-evading by the canonical, “EJC model” NMD rule^5,6,43^ – i.e. PTCs located in the last exon or 55 nucleotides upstream of the last exon junction. We also considered the 2 additional rules implemented in our recent “NMDetective-B” decision tree model^22,44^ – the “start-proximal” rule of NMD escape in the first 150 coding nucleotides, or the partial NMD evasion “long-exon” rule in exons longer than 400nt. Indeed, the NMD-evading PTCs classified by NMDetective-B rules showed lower NMD efficiency across all three variant sets (TCGA somatic mutations: mean log_2_ ratio of 1.67 in NMD-triggering vs. 0.71 in NMD-escaping; TCGA germline variants: 2.27 vs. 0.36; GTEx rare germline variants: 1.15 vs. 0.18; Fig. S2). Encouragingly, while NMDetective-B decision tree was developed using the TCGA somatic mutation dataset^22^ (here we employ an ASE-based quantification approach^21,24^), this data suggests that the two additional, germline PTC datasets have similar or higher signal and therefore utility for developing and/or testing more complex NMD models. Indeed, there is likely a need for these additional datasets, as the NMDetective-B captured about ∼2/3 of the total explainable variance in NMD efficiency acting upon held-out PTC variants^22^. Improving this would not only allow more accurate prediction of PTC pathogenicity in genetic disease and in cancer but has the potential to refine mechanistic knowledge about NMD.

### RNA foundation model used to derive NMDetective-AI, accurate predictor of NMD efficiency genome-wide

We developed NMDetective-AI, a deep learning model that builds upon the Orthrus genomic foundation model for mRNA^28^ finetuned for sequence-based prediction of NMD efficiency. Orthrus is a Mamba-based model with ∼10M parameters pretrained on mature mRNAs i.e. without introns, but with encoded exonic sequence and explicit positions of 5’ UTR, CDS, 5’ UTR and crucially, splice sites, pretrained on ∼45M transcripts from diverse mammalian organisms^28^. Our study fine-tunes Orthrus foundation model on human full-length transcript sequences (including 5’ UTR, CDS, and 3’ UTR) containing PTC variants, to predict the normalized NMD efficiency of a PTC variant (Fig. 1).

For the RNA language model training, we further processed the PTC datasets by more quality filtering (see Methods; Fig. S2-3), correcting for systematic biases (tissue, overlap between variant sets, expression variability), and aggregating recurrent variants by median NMD efficiency, yielding 15,200 unique somatic, and 1,065 unique germline PTCs from TCGA, and 763 unique germline PTCs from GTEx (Fig. 1). NMD efficiency values were then normalized using a linear transformation based on the mean efficiency of NMD-triggering variants (those predicted to undergo NMD by established rules) and last-exon variants (predicted to fully escape NMD), mapping the data to a standardized range (performed on all datasets independently), where +0.5 corresponds to full NMD efficiency upon that PTC and -0.5 corresponds to no NMD efficiency (PTCs in last exons; Fig. S3). This quantity is not strictly bounded, due to measurement noise or prediction noise, actual values can fall outside that range (Fig. S3).

NMDetective-AI was trained on 14,337 somatic premature termination codon (PTC) variants from TCGA, using chromosomes 1 and 20 as a held-out validation set (863 high quality, unique variants). Sequences were encoded as 6-track one-hot representations: 4 nucleotide tracks, 1 coding sequence (CDS) track marking the start of each codon, and 1 splice site track indicating exon-exon junctions (Fig. 1). To identify optimal model configurations, including fine-tuning parameters such as learning rate (see Methods) and the prediction head architecture, we performed a Bayesian hyperparameter search using the W&B tool, while minimizing the validation set loss (Fig. S4) to obtain the final model, henceforth “NMDetective-AI”.

To assess the contribution of pretrained Orthrus representations, versus the Orthrus neural net architecture alone but excluding pretrained information, we compared three training strategies: (1) full fine-tuning of the entire neural net including all layers of the Orthrus encoder, (2) probing with frozen Orthrus encoder weights, training only the added regression head, and (3) random initialization of the Orthrus encoder without pretraining. Training iteration curves revealed stark differences in dynamics and the final performance on validation set loss function across these approaches (Fig. 2a). Full fine-tuning achieved the best performance with validation Spearman ρ=0.668 (R²=0.437, calculated from sum of squares, see Methods), while the limited approach to fine-tuning a head for probing with frozen encoder RNA embeddings reached a maximum ρ=0.408 (R²=0.166), plateauing earlier. This demonstrates that while pretrained features (embeddings) capture useful patterns, task-specific adaptation of the pre-trained neural net optimizes NMD prediction. Remarkably, random initialization without pretraining achieved only ρ=0.068 (R²=0.004) correlation, confirming the utility of the Orthrus as a mRNA foundation model with representations, derived from massive data^28^, that are effectively repurposed when fine-tuning on a modestly-sized dataset such as ours.

**Figure 2:**
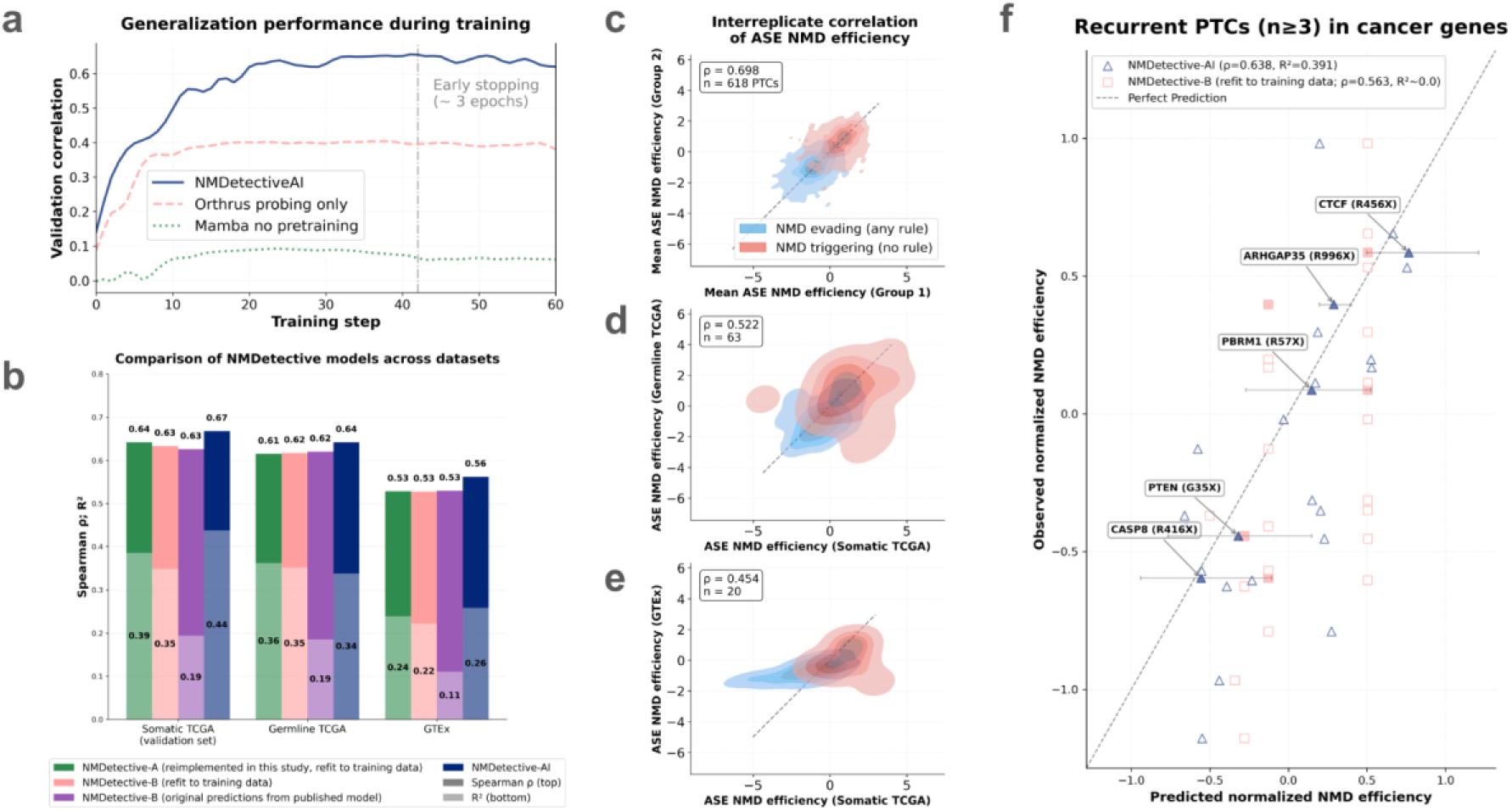
NMDetective-AI model development and validation. (a) Pearson correlation on the validation set during training for three model variants: NMDetective-AI (finetuned Orthrus encoder and regression head), Orthrus probing only (frozen Orthrus encoder with trained regression head), and Mamba no pretraining (randomly initialized encoder with the same configuration as Orthrus). Early stopping occurred at step 42 (approximately 3 epochs). (b) Comparison of NMDetective-AI performance with previous methods across three datasets: somatic TCGA (validation chromosomes only), germline TCGA, and GTEx. NMDetective-A: Random Forest regression model; NMDetective-B (refit to training data): decision tree classifier; these two models were reimplemented in this study following its original design but using the updated datasets. NMDetective-B (original): fixed decision tree with manually defined thresholds as published; NMDetective-AI: this study. Spearman correlation (ρ) and R² values (see Methods) shown. (c) Split-half reliability of ASE NMD efficiency measurements across biological replicates. PTCs with ≥3 observations were randomly split into two groups, mean NMD efficiency calculated per group, and correlation measured across 100 random splits. Red: NMD-triggering PTCs; Blue: NMD-evading PTCs. (d-e) Correlation between ASE NMD efficiency in somatic TCGA and germline TCGA (d) or GTEx (e) for overlapping variants (median values per variant). (f) Scatter plot comparing NMDetective-B (pink square) and NMDetective-AI (blue triangle) model predictions (x-axis) and observed NMD efficiency (y-axis, median across replicates) for recurrent PTCs (≥3 observations) in cancer genes (tumor suppressors and oncogenes). Horizontal error bars on randomly selected variants (filled in symbols; and textual annotations) indicate 25th-75th percentile range across observed replicates. Unselected variants shown in hollow symbols and without error bars to improve visual clarity. Spearman correlation and R² shown for each model. The dashed line indicates perfect agreement in panels (d-g).

We evaluated NMDetective-AI’s ability to generalize to germline variants – crucial for applications to interpreting pathogenic PTC causing genetic disease (Fig. 2b). On germline TCGA variants, the model achieved Spearman ρ=0.642, and R²=0.338. Performance on GTEx ρ=0.562, R²=0.258) suggesting broadly consistent performance in the cancerous somatic tissues and in normal samples. Moreover, NMDetective-AI substantially outperformed our previous generation hand-crafted feature-based approaches, NMDetective-A and B ^22^ across all test sets (Fig. 2b; here we have refitted the NMDetective-A Random Forest model to match our transformed distribution of NMD efficiency labels). On the held-out validation chromosomes from somatic TCGA PTCs, NMDetective-AI achieved Spearman ρ=0.668, and R²=0.437, compared to NMDetective-B’s ρ=0.633, R²=0.348, while NMDetective-A (refit) achieved ρ=0.642, R²=0.385. The performance advantage of NMDetective-AI was consistent across germline TCGA (ρ improvement of 0.642 vs 0.617 for NMDetective-B) and GTEx (ρ of 0.562 vs 0.528 for NMDetective-B), demonstrating that the learned mRNA sequence representations via a foundation model fine-tuned on a targeted, high-quality dataset of causal PTCs has provided a robust, universal NMD predictive model.

To contextualize model performance in terms of maximum attainable accuracy adjusted for measurement error (as before for NMDetective), we assessed the reproducibility of ASE-based NMD efficiency measurements (Fig. 2c-e) across 618 PTCs with ≥3 repeated occurrences. Inter-replicate correlation was Spearman ρ=0.697, establishing an empirical upper bound for prediction accuracy (Fig. 2c). NMDetective-AI approaches this ceiling on the somatic TCGA validation set (ρ=0.668, i.e. 96% of maximum, Fig. 2b). Comparing ASE measurements for the same variants across the somatic data and the 2 germline datasets revealed the inter-replicate correlations in these settings: somatic vs germline TCGA ρ=0.519 (Fig. 2d), and somatic TCGA vs GTEx ρ=0.454 (Fig. 2e). We note that NMDetective-AI performance estimated via Spearman rho on these datasets was slightly above these numbers (see above), suggesting NMDetective-AI models NMD activity upon germline PTCs highly accurately and additionally that the model may be “smoothing over” some technical and biological differences between datasets, which we see as desirable.

To highlight examples of application, we tested NMDetective-AI on 26 recurrently mutated PTCs (≥3 observations, somatic SNVs in the TCGA cohort) in known cancer driver genes (Fig. 2f). In these cases of clinically important somatic PTC mutations, accurate NMD prediction could inform patient stratification and therapeutic targeting by NMD inhibitors (reviewed in^34^ and readthrough agents^37^). NMDetective-AI yielded Spearman ρ=0.638, outperforming our previous NMDetective-B (ρ=0.563). NMDetective-AI predictions of NMD activity upon these 26 recurrent variants were mostly (17/25) within the interquartile range (shown on selected PTCs in Fig. 2f) of the observed NMD efficiency scores (diagonal in Fig. 2f), illustrating the utility of NMDetective-AI for characterizing NMD effects of pathogenic mutations.

### Large-scale mutagenesis to measure NMD in human cells

NMDetective-AI shows exceptional predictive accuracy therefore we reasoned we could use it to systematically re-assess and refine the established NMD rules, considering NMDetective-AI predictions that deviate from binary NMD-escape/trigger classifications, particularly for edge cases near rule boundaries.

Firstly, we considered the classical last-exon/50-nt rule, which has ample prior experimental evidence, yet there is an opportunity to better quantitate the partial NMD evasion near the 50 nt boundary. Secondly, we considered the start-proximal rule of NMD evasion, which has sparse experimental evidence yet in our genomics analyses has fairly prominent coverage of PTCs (12% in NMDetective-B model^22^) therefore needs further study to assess the length of the evading segment, the strength of the evasion as a function of distance from the start codon, as well as the variation of these quantities across genes. Thirdly, we examined the long-exon NMD evasion rule identified in our genomic analyses^22^. However, to our knowledge, this rule lacks direct experimental validation and requires further support, as well as precise quantification of NMD effects across exon lengths and PTC positions within the exon.

Probing the genomics-based NMDetective-AI by model in silico mutagenesis (ISM) can provide insight into the robust patterns observed across training data, thus refining the NMD rules, however we reasoned that controlled experimental testing allows external testing of the rules and the refinements, and offers crucial evidence to support the genomics analysis with controlled perturbation. To this end, we designed various deep mutational scanning (DMS) libraries for measuring degradation, via NMD, of PTC variants introduced into minigene reporter constructs in a human cell line (Fig. 3a). Libraries were ordered as synthesized oligonucleotides or generated with nicking mutagenesis, a primer-based mutagenesis strategy^45^ (see Methods), cloned into a mammalian NMD-reporter and transfected into HeLa cells 48h post-transfection mRNA was extracted, reverse transcribed to cDNA, amplified and sequenced with short-read (Illumina) or long-read (Nanopore, ONT) platforms; referred to as ‘output’. Library DNA was also sequenced before transfection to quantify variant frequency in the absence of selection, referred to as ‘input’. We included non-PTC control variants for normalization: for some libraries, we designed the same set of variants encoding UGG (tryptophan) instead of a PTC; for others, we encoded a non-PTC variant with the original WT sequence. We calculated mRNA level scores and associated errors using DiMSum^46^ (Methods). Briefly, the RNA levels of each variant correspond to the ratio of its frequency after (output) and before (input) selection, normalized to the WT and log transformed (base e). Scores of <0, and ≥0 indicate lower and equal-or-higher RNA levels than the wild-type; respectively. Variants with <10 reads were discarded.

**Figure 3:**
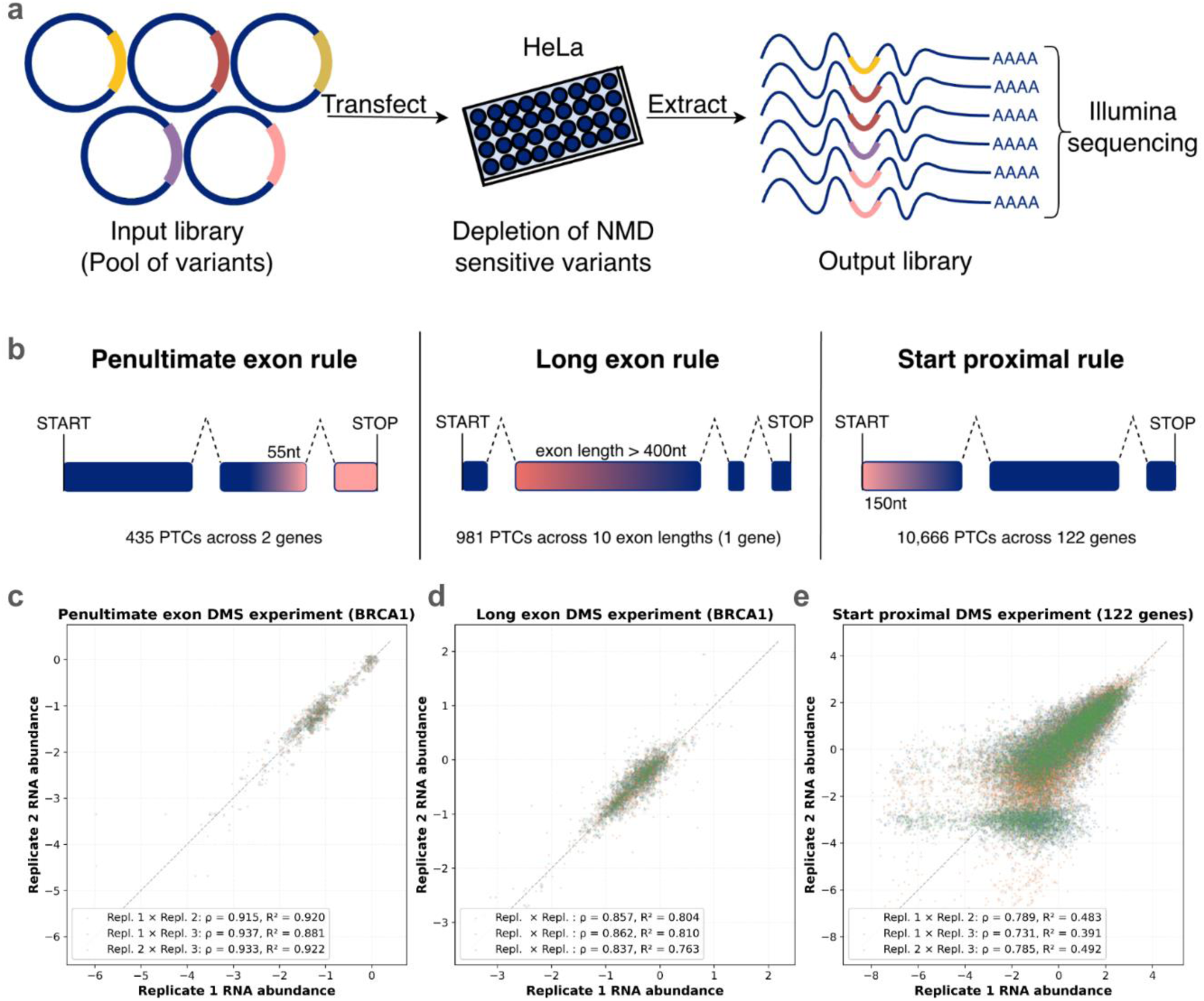
Mutagenesis experiment overview and inter-replicate correlation. (a) Schematic overview of deep mutational scanning (DMS) experimental setup to measure NMD degradation of PTC bearing RNAs. (b) Schematic depicting the NMD evasion rules tested in DMS experiments. (c-e) Inter-replicate correlation for DMS experiments. Grid of scatter plots showing biological replicate comparisons for three DMS datasets (50 nt/penultimate exon experiment in *BRCA1* minigene (c), see Fig. S5a for the *ATP7A* minigene; long exon experiment in *BRCA1* (d); start proximal experiment with data pooled over 122 genes (e)). Each point represents one PTC variant, with RNA level measurements from different replicates compared. All three pairwise comparisons of 3 biological repeats are overlaid in different colors (orange: rep1 vs rep2; blue: rep1 vs rep3; green: rep2 vs rep3). Spearman correlation coefficients (ρ) shown in legends. Dashed black line indicates perfect correlation.

We designed independent DMS libraries and minigenes to explore the last-exon/50 nt, long-exon, and start-proximal rules of NMD evasion (Fig. 3b, Fig. 3c-e, Fig. S5), with high correlations between three biological replicates of Spearman ρ≥ 0.92 (penultimate exon 50nt rule, Fig. 3c), ρ≥0.84 (long-exon, Fig. 3d) and ρ≥0.73 for the start-proximal rule (Fig. 3d).

### Refining the 50-nt rule of NMD escape by genomic model and experimental data

To quantitatively characterize the canonical, last-exon/50-55 nucleotide rule (Fig. 3b), we mutated the 3’-most 76 amino acid positions (228 nucleotides) of the penultimate exons of *BRCA1* and *ATP7A* to all three stop codons (UAG, UAA, UGA) in their respective minigenes (Fig. 4a). We measured RNA abundance for 293 variants per gene (Fig. S6), with Illumina sequencing median coverage of >=190 reads per variant and inter-replicate correlations of ρ≥0.92 (Fig. S5a, Fig. 3c). To match the directionality of our NMD efficiency predictions from NMDetective-AI, RNA abundance levels from DMS were sign-flipped and normalized to the scale of predictions i.e. full NMD efficiency at +0.5 and no NMD at -0.5 (Fig. S3).

**Figure 4:**
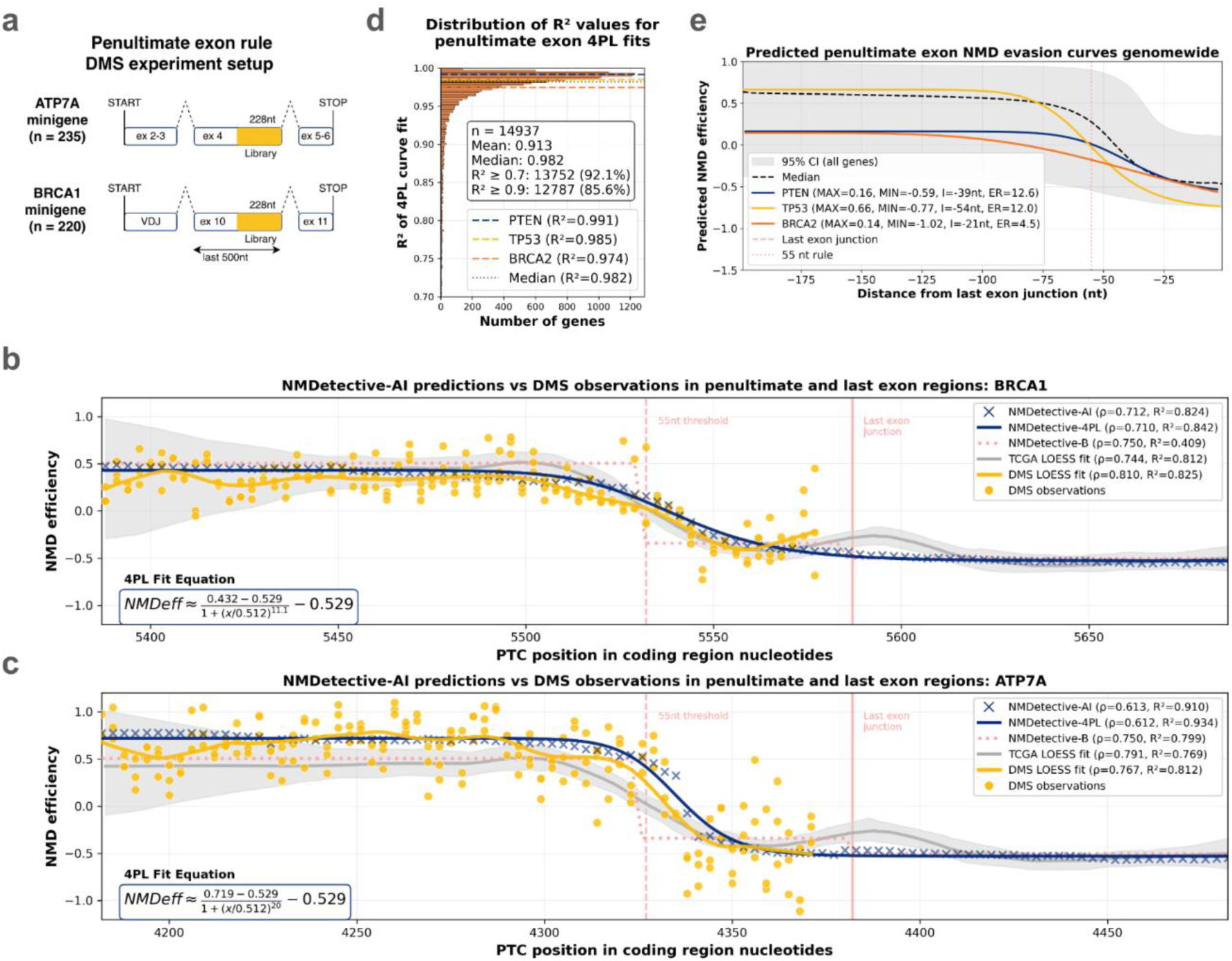
NMDetective-AI captures and predicts quantitative NMD evasion in penultimate and last exons. (a) Schematic of the penultimate exon DMS experiment setup. (b-c) Comparison of NMDetective-AI predictions with DMS observations (all stop types pooled, see Fig. S6 for individual stop type effects) for PTCs in penultimate and last exon regions of BRCA1 (b) and ATP7A (c) genes. NMDetective-AI predictions (crosses) are shown alongside DMS observations (yellow circles with LOESS curve). Background shows LOESS-smoothed NMD efficiency from TCGA PTCs across all genes (gray curve with 95% confidence interval). Vertical lines indicate the last exon junction (solid) and 55-nucleotide rule boundary (dashed). Spearman correlations (ρ) and R² values between NMDetective-AI predictions and DMS LOESS curve are shown in legends. A 4 parameter logistic (4PL) curve, formula in bottom left corners, fit to the predictions are shown in dark blue solid lines. (d) R² values of the 4PL curve fits to genome-wide PTC scan predictions, R² values of selected genes highlighted as dashed lines. (e) Fitted 4PL NMD evasion curves of different genes from PTC scan. MAX: Maximum NMD efficiency, MIN: Minimum NMD efficiency, I: Inflection point, ER: NMD evasion rate. Shaded light-gray band: 95% CI across all penultimate-exon predictions; dashed black line: genome-wide median. Solid colored lines: 4PL curves for PTEN (dark blue), TP53 (gold), BRCA2 (orange) based on NMDetective-AI predictions; vertical dashed: last exon junction, dotted at −55 nt = 55-nt NMD rule.

DMS data revealed a gradual, rather than categorical, change in NMD evasion responses in the exon junction-proximal penultimate exon region (Fig. 4b-c, Fig. S6). LOESS smoothing (R²=0.83 for *BRCA1*; Fig. 4b, and R²=0.81 for *ATP7A*; Fig. 4c) showed NMD efficiency progressively decreased across the -54 to -42 nucleotide window, which meant an RNA abundance increase from ∼30% to ∼100% of the wildtype in both genes. PTCs beyond -54 nucleotides upstream exhibited 2.7-fold and 4.6-fold greater NMD efficiency than those within the -42 nucleotide boundary (*ATP7A*, *BRCA1*, respectively; *p* < 0.01 for both). This gradual transition suggests positional variability in EJC interaction with the NMD machinery.

The DMS experiment data LOESS curve showed strong concordance with LOESS fit to somatic TCGA PTC mutation data in penultimate exon regions across both genes (Fig. 4b-c; n=553 penultimate exon PTCs in TCGA, ρ=0.79 for *ATP7A*, ρ=0.74 for *BRCA1*), validating that experimental minigene systems recapitulate endogenous NMD responses. NMDetective-AI predictions demonstrated strong agreement with both DMS observations, outperforming our previous decision model that outputs a binary response (NMDetective-B, re-implemented here): R²=0.91 vs 0.80 for *ATP7A*, and R²=0.82 vs 0.41 for *BRCA1*. These very high correlations between predictions of NMDetective-AI and the DMS experimental readouts reassure about the quality of both the predictions and the experimental data.

To characterize the NMD transition quantitatively, we fitted four-parameter logistic curves to the NMDetective-AI predictions (see Methods), capturing the gradual decline in NMD efficiency (R²=0.84, and R²=0.93 for ATP7A and BRCA1 gene DMS levels, respectively). The curve had inflection points at nucleotide -51 for both *ATP7A* and *BRCA1* predictions, with local NMD evasion rate indicating smooth transitions (Fig. 4b-c).

We further performed in silico mutagenesis, predicting the NMD efficiency of PTCs at each position across all genes with NMDetective-AI, and fitted four-parameter logistic curves to predictions in penultimate exon regions (Table S1). Analysis of ∼15k transcripts (with ≥3 coding exons; Methods) revealed consistent logistic-shaped transitions (Fig. 4d-e, Fig. S7), with high-quality fits to predictions (R² ≥ 0.7) achieved for 92% of transcripts (Fig. 4d), validating the simple logistic model as an interpretable yet local model of the 50nt penultimate exon NMD escape. The mean NMD efficiency changed from 0.5 (“MAX” in Fig 4e, *sd* across genes=0.34) towards -0.53 (“MIN” in Fig 4e, *sd*=0.27) at the last exon junction. The mean inflection point (“I” in Fig. 4e) occurred at ∼54 nucleotides upstream (*sd*=30.95; Fig. S7). The rate of NMD evasion (ER, Fig. 4e) was similar to *BRCA1* (with a value of 20; Fig. 4b) for the majority of genes (Fig. S7), with a tail of genes exhibiting a more gradual change, like *TP53, PTEN, BRCA2* (Fig. 4e), and *ATP7A* (Fig. 4c). Fitted logistic parameters for the 50-nt rule for all transcripts are available in Table S1.

Overall, this shows that a refined quantitative 50-nt rule of NMD evasion, revised from a step function to a logistic one, is precisely predicted by our genomic language model.

### Long-exon rule of NMD escape fully explored and refined by AI and by systematic experiment

Next, we systematically characterized NMD evasion in long exons, which was discovered in our cancer genomics analyses^6^ and suggested to act at an approximately 400nt exon length. To provide experimental evidence for the long-exon rule and to assess its strength across exon lengths and across PTC positions within the exon, we generated a DMS library spanning nine exon lengths and 82 PTC positions in *BRCA1* exon 10, integrated into a BRCA1 minigene construct (Methods, Fig. 5a, Fig. S8). We created nine 5’-truncated versions (125, 250, 500, 750, 1000, 1500, 2500, 3000, 3426 nt) and introduced stop codons (UAG, UAA, UGA) at all VYY codon positions (V = not T/U, Y = C or T) using nicking mutagenesis^45^, yielding 948 characterized variants. Long-read sequencing achieved high inter-replicate correlation (Fig. 3d) and coverage (median 155 reads per variant). Validation against Illumina-sequenced (from the 50nts rule experiment) overlapping variants (n=21) confirmed assay accuracy (ρ=0.90).

**Figure 5:**
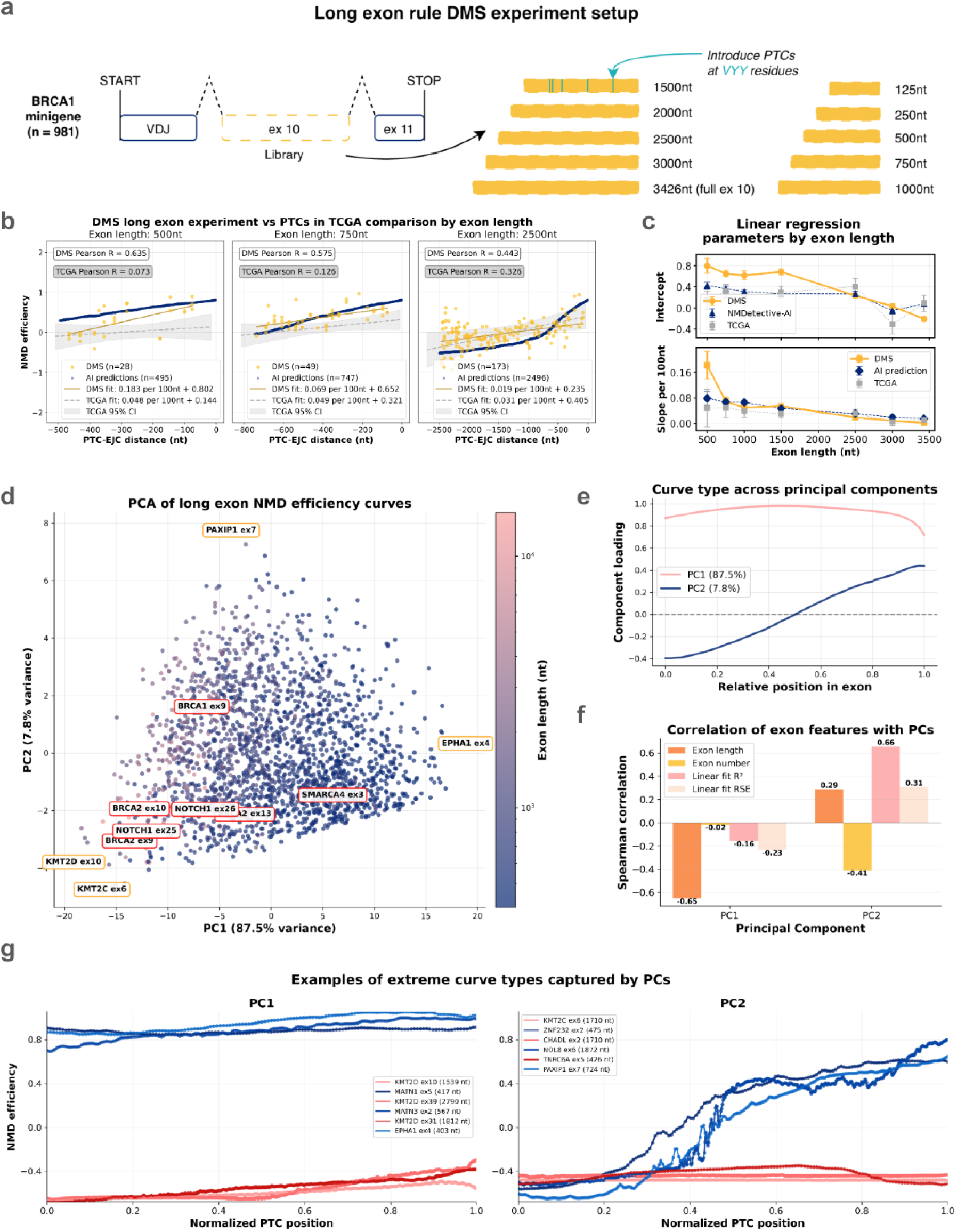
Systematic analysis of exon length and spatial determinants of NMD efficiency. (a) Schematic overview of long exon mutagenesis experiment design (V = not T/U, Y = C or T). (b) Comparison of NMD efficiency between DMS long exon experiment (yellow circles) and TCGA PTCs (gray squares) across three exon length categories (500bp, 750bp, 2500bp). Each panel shows scatter plots with linear regression fits (solid lines) and correlation coefficients. NMDetective-AI predictions shown as crosses. The x-axis shows distance from the exon-exon junction (negative values indicate upstream positions). (c) Linear regressionparameters (intercept and slope per 100nt) plotted against exon length for DMS data (yellow circles), NMDetective-AI predictions (triangles), and TCGA PTCs (gray squares). Error bars show standard errors from the linear models. (d) Principal component analysis of long exon NMD efficiency curves from genome-wide predictions. PCA scatter plot colored by exon length (log scale) with highlighted extreme examples (orange circles) and selected cancer genes (red circles). Variance explained by PC1 and PC2 indicated on axes. (e) Principal component loadings showing NMD efficiency curve patterns captured by PC1-PC2. X-axis shows the relative position within exon (0=start, 1=end of exon). (f) Spearman correlation between exon features (exon length and exon number) and principal components PC1-PC2. (g) Example NMD efficiency curves for exons at extreme positions along PC1 and PC2 axes, showing diverse patterns of NMD evasion across exon positions. The x-axis shows the PTC position normalized between the 5’ end of the exon (x=0) and the 3’ end (x=1).

The mutagenesis data revealed position-dependent NMD evasion for long exons (≥500 nt; Fig. 5b), while shorter exons showed minimal evasion (Fig. S8-10). NMD efficiency increased progressively from the 5’ to 3’ end of exons, consistent with a hypothesized mechanism of the decreased proximity of the stalled ribosome to the EJC (Pearson R=0.47 to 0.65, for 500 to 1500 bp long exons; Fig. S9b-c). However, the magnitude of this PTC positional effect varied substantially by exon length (Fig. 5b, Fig. S8-10). Linear models fit on PTC position demonstrated that longer exons exhibited weaker positional gradients (smaller slope coefficients) but a higher baseline NMD evasion (decreased intercepts meaning overall low NMD efficiency irrespective of position). For exons ≥500 nt, regression slopes decreased consistently from 0.18 increase in NMD efficiency (on our scale of -0.5 to +0.5) per 100 nucleotides (at 500nt) to only 0.02 increase per 100 nucleotides (at 3426nt). The intercept values (Fig. 5c) - the NMD efficiency at the 3’ end of the exon - decreased from 0.8 to -0.2; thus, in very long exons, NMD escape is strong and fairly independent of PTC position. This effect can also be observed when comparing the same PTC-to-downstream splice site distance across exon lengths, which shows a strong exon length effect on NMD, independent of the PTC-to-downstream splice site distance (Fig. S9a).

The experimental data also demonstrated agreement with somatic TCGA PTC mutation RNA-Seq ASE signal in matched exon length categories pooled across different transcripts (Fig. 5b, Fig. S10). The linear fit to genomic PTCs was an adequate model for DMS measurements in exon categories with enough data. For 500 nt and 750 nt exons, DMS measurements and the linear fit to local NMD efficiency of TCGA mutations correlated at R=0.65 and 0.62, respectively. For extremely long 2500 nt exons, agreement was R=0.38, likely in part due to sparse sampling of very long exons in TCGA mutations (119, 163, and 55 PTCs for exon length categories 500nt, 750nt, 2500nt respectively). The characteristics of TCGA linear models showed similar trends of decreasing slope and intercept values as DMS data (Fig. 5c).

To achieve comprehensive PTC coverage at nucleotide-resolution, we performed in silico PTC scanning with NMDetective-AI across all *BRCA1* exon length variants. Predictions captured the gradual 5’-to-3’ NMD efficiency increase observed in the experimental data, while filling the gaps between sparse DMS observations (Fig. 5b, Fig. S10). Linear fits to the predictions also exhibited the similar trends across exon length categories up to 1500 nt (Fig. 5c). However, in very long exons (≥2500 bp) exhibited a piecewise NMD efficiency profile (Fig. 5b, Fig. S7): NMDetective-AI predicted NMD efficiency increases gradually towards the 3’ end of exons, with a sudden increase in the last ∼1000 nts of the exon. Thus, the genomic data suggests that in extremely long exons, the linear fit of NMD evasion efficiency to the PTC coordinate works only within the 5’-most 1500 nt, after which NMD evasion dwindles.

Finally, we asked if there exist systematic differences in the long exon NMD patterns across the individual genes/exons (Fig. S11). In a genome-wide NMDetective-AI run, we extended in silico scanning to 2,171 exons with lengths >400 nt, excluding the first and last two coding exons, which have separate NMD-escape mechanisms. Principal component analysis on NMD efficiency curves (50 uniform positions sampled per exon, with interpolation) revealed variation in long-exon NMD patterns (Fig. 5d). PC1 explained 86% variance and captured the overall magnitude of NMD evasion (Fig. 5e), strongly associating with exon length (Spearman ρ=-0.52; Fig. 5f). PC2 (7.8% variance) distinguished between linear-like gradients and plateau-like profiles where NMD efficiency saturates in the 3’ exon region (Fig. 5e), suggesting that some long-exons -- those with extreme PC2 values -- might exhibit some systematic differences in NMD for features apart from length itself. Interestingly, PC2 correlated with the exon order within the transcript (ρ=0.41; Fig. 5f), suggesting that exons closer to the 3’ of transcripts exhibit flatter NMD profiles. The PC3 explained considerably less signal (3.5% variance) and so we do not further interpret the PC3 or lower-ranking PCs. NMD profiles of exons with highest or lowest PC2 values are shown on Fig. 5g, with further example long exons shown in the supplementary (Fig. S11) as well as full PC data provided in Table S2.

### Start-proximal NMD escape rule systematically assessed by experiment exhibits gene-level variation

After comprehensively characterizing the long-exon NMD evasion, we next turned to address the start-proximal NMD evasion rule by combination of experiment and genomic language model analyses (Fig. 3b). To study this rule across diverse genetic contexts, we performed DMS on 139 genes with high pathogenic nonsense mutation burden, selected from ClinVar for genetic disease, and MSK-IMPACT and TCGA databases for somatic PTCs (Fig. 6a). For each gene, we designed the non-stop wild-type and the sequence variants with a UAG stop codon at every position within the 5’-most 83 amino acids. The library was cloned in a *BRCA1* minigene reporter containing exons 1-11 and three introns (Fig. 6a). After filtering for reproducibility (≥10 reads, mean DimSum error estimate σ>1 across three biological replicates), 10,066 variants over 122 genes were retained (91% of library) with good inter-replicate correlations (ρ≥0.73, Fig. 3e). We further retained only the genes where majority of PTCs exhibited expression signal equal to or lower than the wild-type signal (n=100, see Methods for all filters, Fig. S12-S13), and processed observations to harmonize with the scale of our NMD efficiency labels (see Methods). Namely, the NMD efficiency was normalized to be highest (i.e. no evasion; corresponding to +0.5 on our scale) by considering the 3’ end of every gene, under the reasonable assumption that start-proximal NMD evasion ceases after approximately 250 nt. Furthermore, based on pooled genomic PTCs (Fig. 6b) we observed that the 5’ end evasion is total (corresponding to -0.5 on our scale) and thus rescaled the DMS data 5’ end NMD efficiency globally to match that, while maintaining possible inter-gene heterogeneity in the degree of 5’ end evasion (see Methods). To ensure that the minigene setup did not cause aberrant splicing and mask nonsense-mediated decay (NMD) effects, we utilized SpliceAI^47^ to predict potential alternative splice sites within our library. Our analysis demonstrated that predicted missplicing was not associated with NMD resistance, confirming that our results were not confounded by unstable isoforms (Fig. S15d).

**Figure 6:**
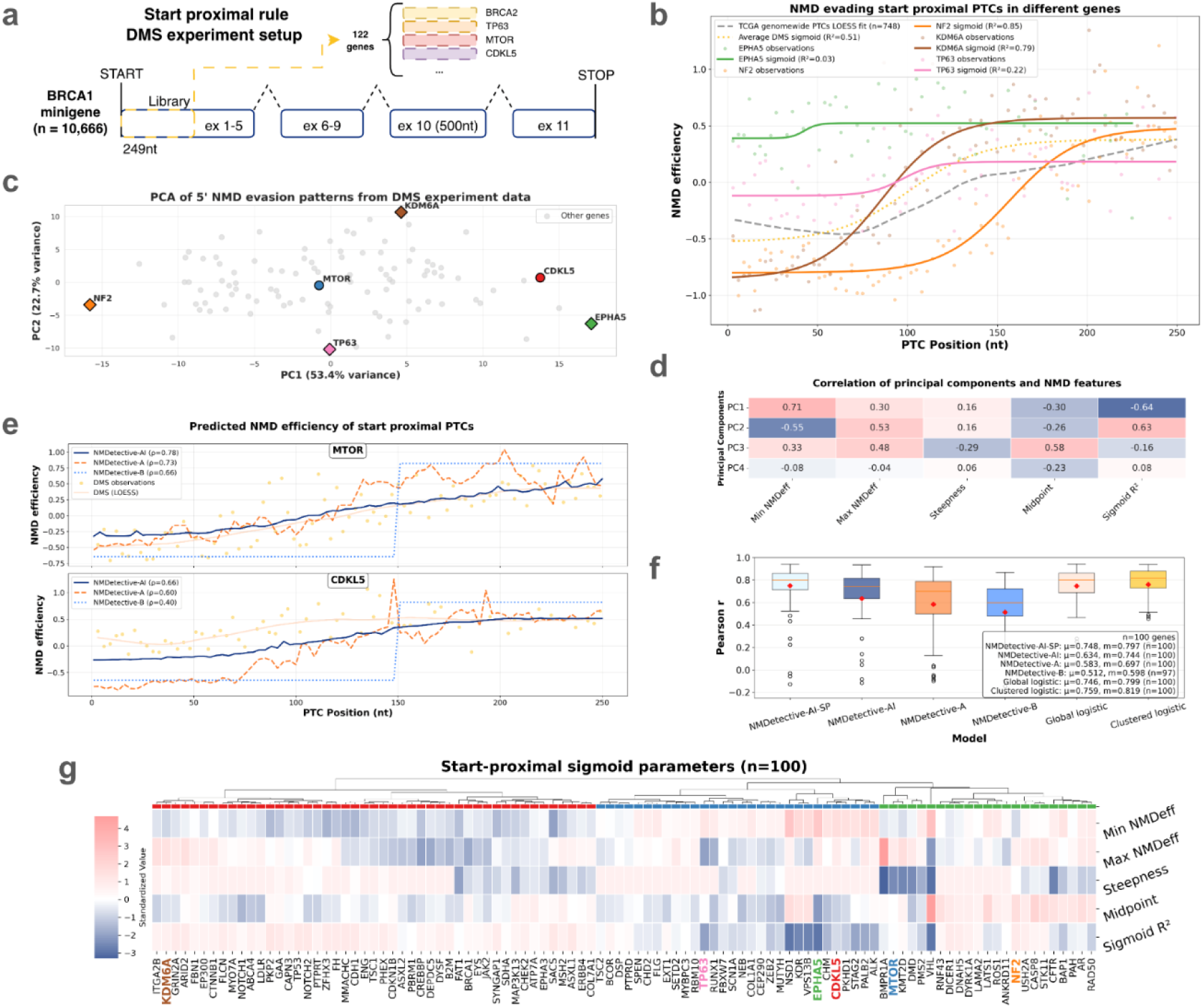
Landscape and predictability of start-proximal NMD evasion. (a) Schematic overview of start-proximal mutagenesis experiment design. (b) NMD efficiency profiles for genes highlighted in panel (c) as squares in the start-proximal region. DMS observations shown as circles with 4-parameter sigmoid curve fits (solid lines). Gray dashed line shows TCGA genome-wide LOESS fit for context. X-axis shows position in coding sequence (nucleotides from start codon). (c) Principal component analysis of start-proximal NMD efficiency curves. PC1 vs PC2 scatter plot with selected genes highlighted (CDKL5, MTOR as colored circles; extreme genes along each axis labeled) and other genes shown in gray. LOESS-interpolated fitness values used for PCA. Variance explained by each PC indicated on axes. (d) Correlation heatmap showing Spearman correlations between principal components (PC1–PC4) and logistic fit parameters (Min NMDeff, Max NMDeff, Steepness, Midpoint) and fit quality (R²). Color scale indicates Spearman correlation coefficient. (e) Predicted NMD efficiency profiles for MTOR and CDKL5 genes. Predictions by NMDetective-AI (blue line), NMDetective-A (dashed orange line), and NMDetective-B (dotted light blue line); DMS observations (yellow circles and LOESS curve). (f) Model performance comparison across all 100 DMS start-proximal genes. Boxplots show distribution of per-gene Pearson correlations between model predictions and DMS observations for NMDetective-AI-SP, NMDetective-AI, NMDetective-A, NMDetective-B, global sigmoid fit, and cluster-specific sigmoid fit. Sigmoid models evaluated using the same 10-fold gene-level cross-validation framework as NMDetective-AI-SP. (g) Hierarchical clustering heatmap of genes based on standardized 4-parameter sigmoid fit parameters and fit quality. Genes clustered by Ward’s method into 3 groups with cluster assignments shown by color bar.

The experimental data indeed validated the broad relevance of the start-proximal NMD evasion rule in this controlled experiment, where most genes showed NMD activity decreases at the 5’ end (Fig. S12). However, casual inspection also revealed gene-to-gene heterogeneity in start-proximal NMD evasion profiles (Fig. 6b, Fig. S12-13), where magnitude and positional features varied. Four-parameter sigmoid curves (see Methods) provided adequate fits (R²≥0.5) for 80 of 100 genes (median R²=0.71), and their parameters showed wide gene-level variation: the 5’ NMD efficiency (“Min NMDeff”) ranged from −1.23 to 0.45 (median −0.63), and the “Midpoint relative position varied from 0.15 to 1.00 in relative position (median 0.40, or about 100 nts). Moving from experiment to observational, genomic data analysis, LOESS fits to start-proximal PTCs from TCGA (n=748, here pooled across genes) also showed similar sigmoid-like NMD efficiency curves; however, sample size did not permit gene-specific analysis (Fig. 6b).

Next, we aimed to summarize the main trends in heterogeneity between genes in start-proximal NMD escape. A principal component analysis on LOESS-interpolated NMD efficiency curves (83 positions × 100 genes) (Fig. 6c) yielded a PC1 (53% variance) that primarily captured overall NMD evasion magnitude, correlating with 5’-most PTC NMD efficiency (Min NMDeff, Pearson R=0.71). Out of the variables we surveyed, the PC2 (23% variance) correlated most strongly with the fit quality R² (r=0.63), thereby distinguishing genes where the sigmoid model was inadequate (n=20 genes with R²<0.5) from those well-described by it (n=80 genes with R²≥0.5); we do note that the PC1 and PC2 did also correlate to lesser extents with other potentially explanatory variables (see Fig. 6d). PC3 (13% variance) encapsulated the length of the 5’ NMD-evading region, correlating with the Midpoint parameter (r=0.58) most strongly of the variables assessed, separating start-proximal regions of genes with more 5’ versus more 3’ transitions towards the default, NMD triggering state. These dominant axes of systematic variation in start-proximal NMD evasion correspond to interpretable features of the sigmoid response model (Fig. 6d); conceivably, these may have specific biophysical reasons rooted in variable efficiency and/or localization of translation reinitiation underlying the start-proximal NMD escape.

### Start-proximal NMD evasion quantified genome-wide via computational analysis

The start-proximal NMD rule was clearly observed in trends extracted from genome analysis: NMDetective-AI predictions demonstrated strong agreement with DMS observations for individual genes (Fig. 6e). Evaluating model performance systematically across all 100 DMS genes (Fig. 6f, Fig. S12), NMDetective-AI achieved median per-gene Pearson R=0.744, outperforming our previous rule-based model NMDetective-B, which relies on a binary classification rather than local gradients in NMD escape (median R=0.598), as well as NMDetective-A (Fig. 6f).

Next, we explored whether our genomic, sequence-to-function model can be further improved by jointly considering DMS data. We fine-tuned NMDetective-AI on the DMS dataset (appropriately normalized) using 10-fold cross-validation, for each individual gene level assayed in DMS. The specialized model (NMDetective-AI-SP) showed further improvement in median per-gene correlation (R=0.797 vs R=0.744 for base model). Thus, the experimentally observed start-proximal NMD evasion patterns contain gene-specific signals not fully captured by fine-tuning Orthrus on current data, possibly because of noise in training data and/or limitations of mRNA sequence representation in the Orthrus neural net architecture. This gene-specific signal, however, appears quantitatively modest: a global sigmoid-fit (using median parameters across all training genes) achieved median R=0.799, while using multiple sigmoids (based on hierarchical grouping of genes by local evasion patterns) performed modestly better (median R=0.819).

This signal is sufficiently robust that hierarchical clustering of sigmoid parameters (Midpoint, Steepness, Min and Max NMDeff) can be used to identify three gene categories based on evasion patterns (Fig. 6g; all individual gene plots shown in the Fig. S12-S13). Cluster 1 (n=47 genes, e.g. *KDM6A, BRCA1, TP53)* has strong 5’ NMD evasion (median Min NMDeff −0.78), high Steepness, and Midpoints in the first half of the profiled region (median relative coordinate 0.38 (in a theoretical range 0-1)), representing genes with a sharp and pronounced transition from NMD evasion to NMD triggering. Cluster 2 (n=30 genes, e.g. *EPHA5*, *CDKL5*, *TP63*) showed weaker sigmoid fits (median R²=0.49) and a shallow 5’ evasion patterns (median Min NMDeff −0.26), indicating genes where start-proximal NMD evasion is less pronounced and/or does not follow a clear sigmoid pattern. Cluster 3 (n=23 genes, e.g. *DICER1*, *NF2*, *MTOR*, *STK11*, *VHL*) was characterized by Midpoints somewhat shifted away from 5’ end (median relative coordinate 0.55, range 0.26–1.00) and lower Steepness, representing genes with NMD evasion that extends further into the coding sequence with a more gradual transition.

Extending predictions genome-wide using NMDetective-AI on 11,321 protein-coding genes (see Methods for filters), we classified genes again into three shape categories from the parameters of the sigmoid curves fit to predictions (Table S3). The majority of the genes (10,142) exhibited typical sigmoid patterns (Fig. S14), with high R² (0.98) and characteristics similar to that of Cluster 1 above (Min NMDeff -0.24, Max NMDeff 0.56, high Steepness, Midpoint 0.53). The second group (by numerosity, n=925) showed atypical sigmoids, with early Midpoints (0.10), and a continued linear increase of NMD efficiency after the upper plateau was reached (Fig. S14). The third group of genes (254) showed little NMD activation in the first 250nt (Fig. S14). We note that essential and disease-associated genes were depleted in this last category OR=0.39 (*p*=0.049) and OR=0.63 (*p*=0.002), respectively.

### Translation reinitiation promotes NMD evasion at start-proximal gene ends

As an additional component of our DMS library, we designed variants to systematically test the mechanistic hypothesis that translation reinitiation contributes to NMD escape at 5’ gene ends. Using the *BRCA1* start-proximal gene sequence (previously already used as a start-proximal reporter^48^), we designed libraries by applying PTC saturation mutagenesis over the 5’-most 83 residues, where each residue was mutated to the three stop types (Fig. 6a, Fig. 7a). Next, we selected all UGA PTC variants, and, for each, we designed variants by placing AUG start codons at different distances downstream (Fig. 7a). Measuring NMD for each UGA PTC with or without an AUG methionine downstream (referred to as “AUG” and “non-AUG” respectively) enabled us to decouple the effects on NMD evasion due to proximity to the translation start site (TSS) proximity from translation reinitiation effects. The AUGs were introduced at a distance of 1, 5, 10, 20, 30, 40, 50, 60, 70 or 80 residues downstream of the PTC, provided that they were within the 83 residues length of the oligo (Fig. S15a-b). Note that all AUGs from the constant part of the reporter (except the first AUG) were replaced by CUG, so that AUG-driven evasion from NMD could be attributed only to the library-encoded AUGs. In total we measured 1185 variants, with a median coverage of 135 reads per variant and highly correlated across replicates (R=0.84-0.90), using Illumina sequencing.

**Figure 7:**
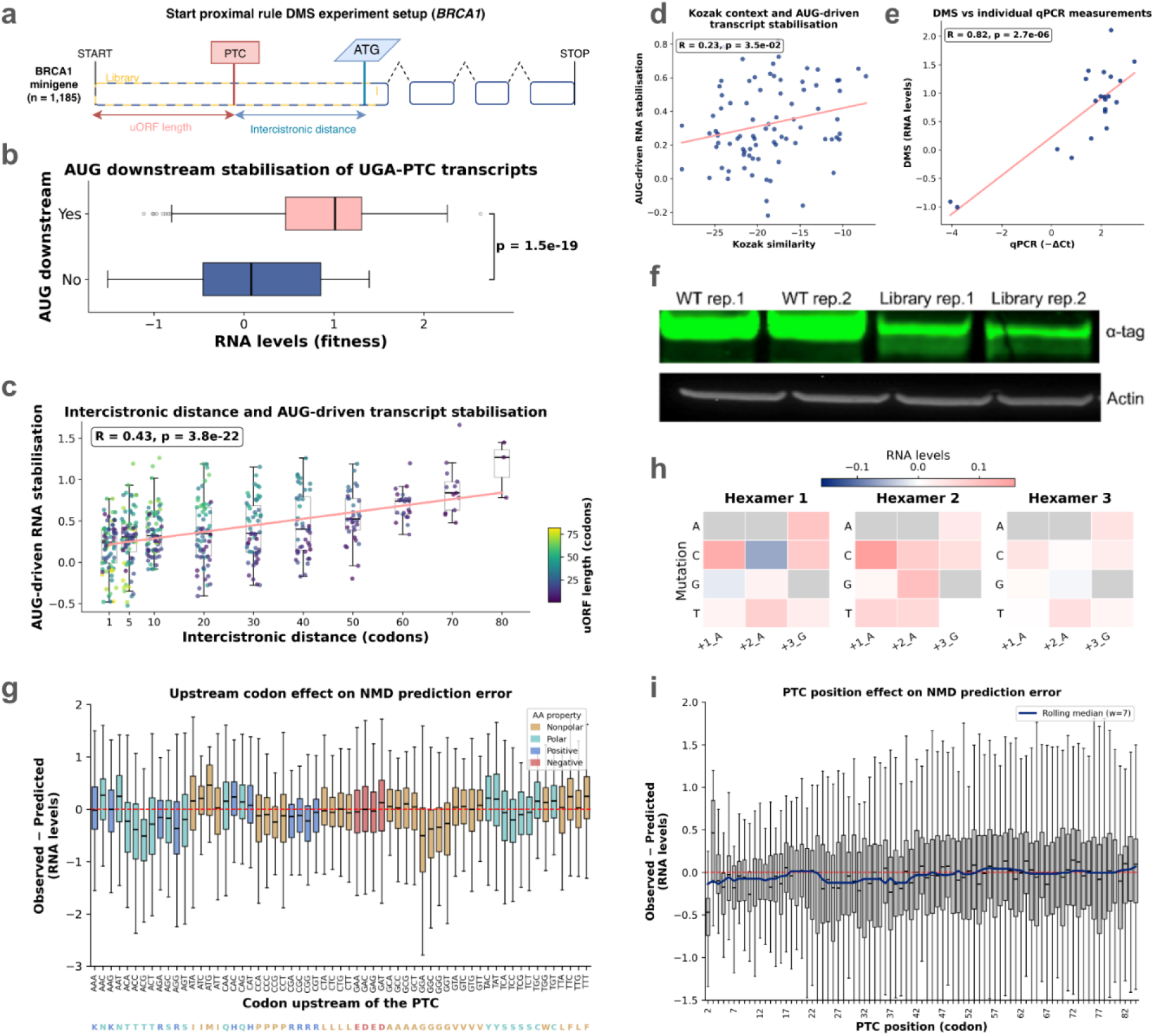
Reinitiation and local sequence-context effects. (a) Library design showing the transcript features explored in the library. (b) Horizontal boxplot comparing RNA levels between UGA-PTC variants with and without a downstream AUG codon; statistical comparison by two-sided t-test. (c) AUG-driven RNA stabilization (increase in fitness relative to the matched AUG-free variant at the same PTC position) as a function of the intercistronic distance (codons between PTC and downstream AUG). Boxplots at each tested intercistronic distance are overlaid with individual data points colored by uORF length (PTC position). A linear regression line is shown with Pearson r and *p*-value. For intercistronic distances ≥10, longer uORFs drive greater stabilization than shorter ones at the same intercistronic distance. (d) Scatter plot of mean AUG-driven RNA stabilization per AUG position plotted against the Kozak sequence similarity score for each position along the *BRCA1* 5′ region. Each point represents one unique AUG position; linear regression shown with Pearson r and P-value. (e) Scatter plot comparing DMS-measured RNA levels with individual qPCR measurements (−ΔCt) for 22 variants spanning the full NMD efficiency range. Pearson r and P-value shown. (f) Western Blot of cells transfected with the WT or the library. The smear below the full-length protein band, detected in the library but not in the WT samples, belongs to N-terminal truncated isoforms arising from translation reinitiation downstream of PTCs. (g) Residuals grouped by the codon immediately upstream of the PTC; boxes colored by the biochemical property of the encoded amino acid (nonpolar, polar, positively charged, negatively charged), with single-letter amino acid annotations below each codon label. (h) Effect of mutations at downstream hexamer positions (+1, +2, +3) on RNA levels. Colors represent regression coefficients: red indicates higher RNA levels (more NMD), blue indicates lower RNA levels (less NMD). (i) Residuals grouped by PTC position (codon number); box width scales with the square root of the number of observations per position, and a rolling-median trend line highlights the overall positional effect. In both panels, the red dashed line at y = 0 indicates no prediction error; negative values indicate over-estimation and positive values indicate under-estimation of RNA levels.

Overall, the presence of AUGs downstream consistently increases RNA abundance compared to non-AUG variants (2.0-fold, *p*<4e-13), suggesting a robust RNA stabilizing effect of translation reinitiation (Fig. 7b). For each PTC, the ‘AUG-driven RNA stabilization’ metric quantifies the RNA levels increase of the AUG variant compared to their non-AUG counterpart (both have the PTC at the same position), hence reporting the AUG effect decoupled from the distance-to-TSS effect. The metric shows significant correlation with two features: distance from the PTC to the downstream AUG (intercistronic distance) and PTC position relative to TSS (upstream ORF length) (Fig. 7c, Fig. S15b). For variants with the PTC in the same position, longer intercistronic distances drive higher RNA levels, suggesting a stabilizing effect (R=0.37, *p*<2e-16). After controlling for the intercistronic distance, transcripts with longer uORF lengths have higher RNA levels (R=0.51, *p*<2e-16) (Fig. 7c). This is observed for intercistronic distances >=10, since shorter intercistronic distances cause very small stabilization which likely masks the uORF length effect. Additionally, the similarity of the AUG-proximal region to the consensus Kozak motif positively correlates with RNA abundance (R=0.23, *p*<3.5e-02), consistent with reinitiation of translation being the stabilizing mechanism, presumably via NMD evasion (Fig. 3d).

We designed the reporter to express an alpha tag encoded at the 3’ end of the transcripts, which was used to validate by Western Blot the occurrence of translation reinitiation. We detected the presence of shorter protein isoforms arising from translation reinitiation events in cells transfected with the full library, which are absent in control cells transfected only with the WT (Fig. 3f). Finally, to validate the DMS results, we individually qPCR-quantified the RNA abundance of 22 *BRCA1* variants spanning the whole dynamic range, obtaining a high correlation (r=0.83) (Fig. 3e).

In summary, our results strongly support translation reinitiation as a mechanism of start-proximal NMD evasion and further suggest that it is modulated by uORF length, intercistronic distance, and the Kozak strength of the AUG-surrounding sequence

### Local sequence context effects on NMD

DMS libraries showed several cases in which NMD levels differed significantly across the three stop types. For instance, in the BRCA1 50nt rule library, we detected a stronger NMD for UAG in four positions (PTCs in -51/-49, -57/-55, -66/-64 and -141/-139 nts, adjusted *p* < 4e-4), some of them with particularly pronounced effects. Other PTCs showed different patterns, including cases where UAA drove either the highest or the lowest NMD, UGA drove the lowest NMD, or where no significant differences were observed among the three stop codons (Fig. S6, Fig. S8-S9, S15a). The lack of consistent stop type effects across positions, suggests that these are shaped by the local sequence.

Consistent with this, we observed a recurrent trend across all DMS datasets where contiguous positions show RNA abundance differences consistent with effects of local context (Fig. S6, Fig. S8-S9, S15a). Our start-proximal DMS library contains a considerable number of data points, thus we used it to explore the effects of the PTC-neighboring nucleotides on NMD efficiency. For each variant (those that RNA levels lower than wildtype of the same gene) we calculated the residuals after regressing out the effects of the PTC position (from a LOESS fit) and tested its association with the sequence upstream and downstream of the PTC. The downstream trinucleotide showed minor differences (Fig. S15c), whereas the upstream codon showed stronger effects, which were largely clustered by the encoded amino acid (Fig. 7g). The amino acids G (1.6-fold decreased RNA levels relative to expected, adjusted *p* < 2.2e-16) and T (1.5-fold, adjusted *p* < 2.2e-16) represented the strongest NMD-sensitive context, whereas amino acids N and F (1.3- and 1.2-fold increased RNA levels relative to expected; respectively, adjusted *p* < 3e-3) drive the highest NMD evasion; the methionines were not assessed for protective effect. This is in overall agreement with a recent study suggesting translation dynamics govern NMD, highlighting the NMD-promoting effect of amino acid G (glycine) preceding the PTC, with T (threonine) showing similar effect^49^. Effects were similar across codons, suggesting a largely amino acid-driven mechanism.

Motivated by these observations, we designed a library to specifically assess the sequence NMD effect of the codon upstream and downstream the PTC. We fully randomized hexamers at three different exon 10 positions of our *BRCA1* reporter (Fig. 6a). Experiments were performed in triplicates (correlations r=0.82-0.9). For each of the 64 codons, we had variants with a PTC upstream or downstream as well as control variants without PTCs, used to control for the NMD-independent effect on RNA stability of each codon. We fitted linear models to predict RNA abundance based on the codon identity for PTCs (and for non-PTC variants used for normalization), separately for the 3 stop types. The normalized coefficients capture the NMD-dependent effect on RNA abundance of the codons downstream and upstream of the PTC. For UAG and UAA stops we did not detect consistent effects across any of the three randomized hexamers, but we observed that +1/+3 CUA trinucleotide protects UGA transcripts from NMD (Fig. 7h). Out of the 64 codons placed downstream of UGA PTCs, CUA is the top-ranked, top-ranked and top four most NMD-resistant contexts across the 3 randomized hexamers. Readthrough over stop codons inhibits NMD^15,37,50^, and UGACUA is the most readthrough sensitive context in humans^51–53^, suggesting that the observed UGACUA NMD escape is due to stop codon readthrough. No upstream codon effects were consistently detected for any stop type.

Lastly, we detected two local features determining RNA levels, shared across genes. First, PTCs at position 1 (which prevent translation initiation by destroying the canonical AUG) display lower RNA abundance than predicted by the PTC position (LOESS fit) (1.5-fold, *p* < 2.2e-16, Fig. 7i), supporting the association between translation and mRNA stability. Second, PTCs at position 2 drive higher RNA levels than predicted via LOESS (1.6-fold, *p* < 2.2e-16) (Fig. 7h). The effects are strong, and both are the positions with the highest observed *vs* predicted error across genes (Fig. 7i). Our data does not distinguish whether the NMD-protective effects of PTCs at position 2 are due to the PTC position or possibly to the methionine codon upstream of the PTC, since we removed the other methionines to prevent translation reinitiation.

### NMD modulates selection on PTC variants in genetic disease and cancer

To assess how NMD shapes the landscape of pathogenicity of stop-gained variants, we analyzed 108,258 rare germline PTCs (allele frequency <0.1% across all gnomAD v4.1 subpopulations) and subsequently 102,598 somatic PTCs from TCGA whole-genome sequencing. Both sets contain a subset of pathogenic PTC variants, in case of gnomAD causing genetic disease with recessive mode of inheritance (since the database contains individuals without overt phenotypes of genetic disease, dominant acting PTCs are less likely to be represented), and in case of TCGA a set of cancer driver somatic PTCs. We assessed the patterns of selection on PTCs in light of their predicted NMD activity -- negative selection on germline PTCs in ostensibly healthy individuals, and positive selection on somatic PTCs in tumor genomes -- to infer if NMD is protective or NMD is aggravating, when it acts upon pathogenic PTCs in a particular gene. Individual examples of both cases were demonstrated (reviewed in^33,35,54^) and our prior genomics analysis on ClinVar^22^ suggested that the NMD aggravating case may be fairly common across genetic diseases, suggesting opportunities for NMD inhibition therapies. Here, we analyze a large population genomics database for additional power and moreover apply NMDetective-AI for improved accuracy of NMD annotations, to infer selection signatures and prioritize disease genes where NMD activity is likely to be pathogenic.

Germline variants were classified using both established NMD evasion rules and NMDetective-AI predictions with thresholds ≤−0.17 for evading, ≥0.43 for triggering, with intermediate predictions excluded in this analysis (see Methods, Fig. 8a, and Fig. S16). Rule-based classification categorized 47.0% rare gnomAD PTCs as NMD-evading (Fig. 8b), among which (Fig. 8c), the largest category was last exon (25.3%), followed by long exon (10.1%), start-proximal (8.2%), and 55-nucleotide penultimate exon rule (3.4%). NMDetective-AI predictions yielded a three-way classification (based on a mixture model of predicted probabilities, Fig. 8b): 46.7% triggering, 32.6% evading, and 20.6% intermediate. Comparing AI predictions with rule-based categories, NMDetective-AI confirmed the majority of last exon variants as evading but classified only some 55-nucleotide rule and long exon variants as crossing the threshold for the evading category, indicating that classical rules may oversimplify interpretation for some variants (Fig. 8c). An additional 729 PTCs classified as NMD-triggering by rules were predicted as evading by NMDetective-AI, representing variants whose evasion mechanism is not captured by established rules and/or usage of alternative transcripts. We note that 208 of these evading cases were located in penultimate exons (outside the final 55 nt thereof), which we noted previously^22^ with unclear mechanism, speculatively relating to incomplete occupancy of exon-junction complexes on splice sites.

**Figure 8:**
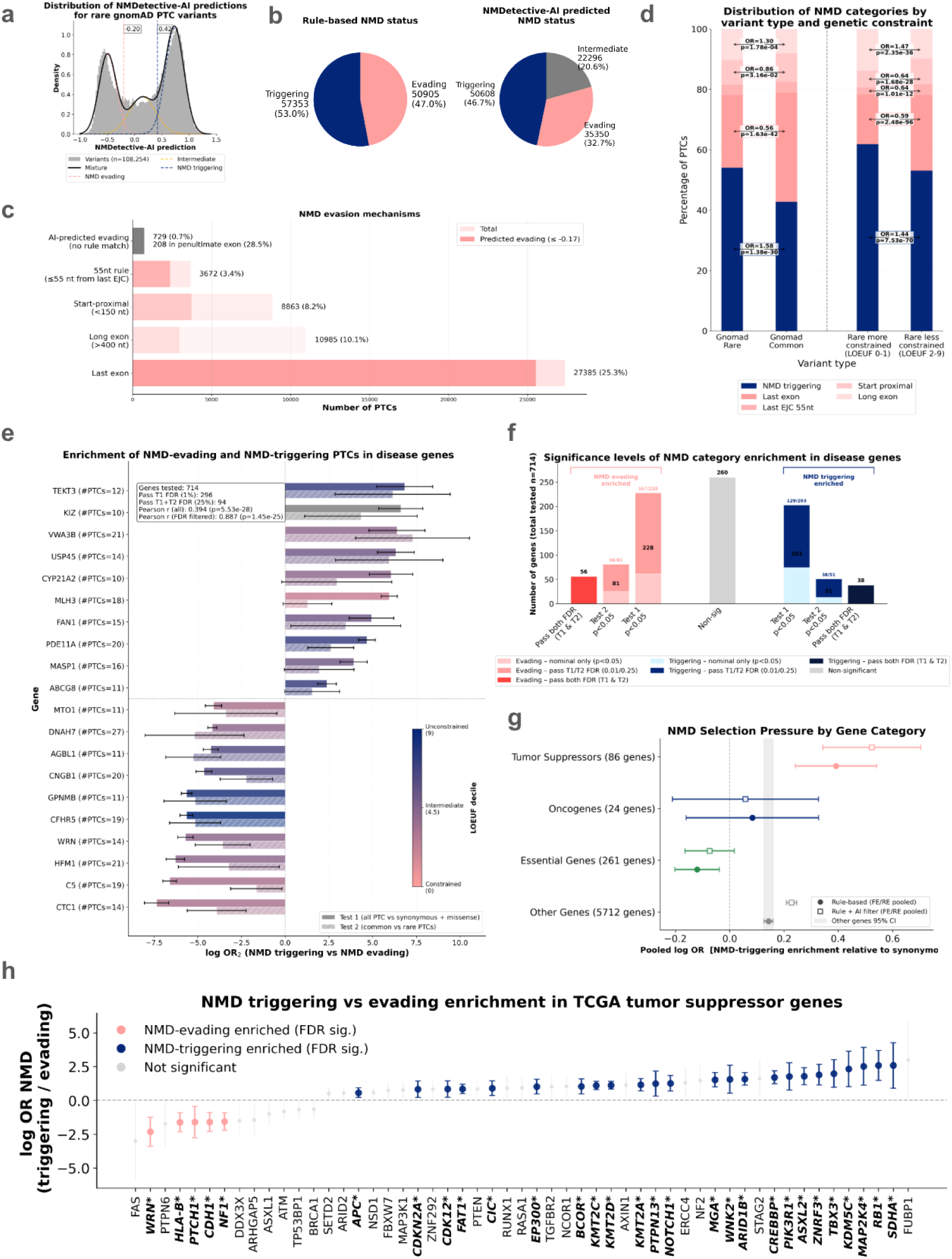
Selection of PTCs in germline and soma. (a) Distribution of NMDetective-AI predictions for rare gnomAD stop-gained variants. A 3-component Gaussian mixture model (GMM) is fitted to the predictions; intersection points between adjacent components define thresholds separating NMD-evading, intermediate, and NMD-triggering categories (dashed vertical lines). (b) The proportion of NMD-triggering and NMD-evading rare PTCs based on established NMD evasion rules (left; for rules see panel c) or NMDetective-AI predictions (right). PTCs in the intermediate category fell between the GMM-derived thresholds (see Methods). (c) Breakdown of NMD evasion mechanisms among rare evading PTCs. Categories: start-proximal (<150 nt from start codon), last exon, 55nt rule (≤55 nt from last exon-exon junction), and long exon (>400 nt). Darker regions show the proportion confirmed by NMDetective-AI predictions. (d) Comparing the distribution of NMD categories between variant types: all rare PTCs, common, and rare PTCs stratified by genetic constraint (LOEUF decile 0–1 vs 2–9). Fisher’s exact test results shown for statistically significant differences between NMD groups (shown where *p* < 0.05). (e) Per-gene (Wald) log odds ratio (with 95% CI) of NMD-triggering to NMD-evading allele counts for the top ClinVar disease genes most enriched in each direction. For each gene, two bars are shown: Test 1 (solid; rare PTCs normalized by synonymous and missense allele counts; see Methods) and Test 2 (hatched; rare vs common within-gene PTC split). Top 10 triggering-enriched and top 10 evading-enriched genes selected by combined *p* value. Positive log OR indicates enrichment for NMD-triggering PTCs; negative for NMD-evading PTCs. (f) Summary of NMD variant enrichment significance for ClinVar disease genes, separated by direction. Enrichment was assessed with two complementary Wald log OR tests (see Methods): Test 1 (T1) compares rare PTC allele counts normalized by synonymous allele counts; Test 2 (T2) uses a within-gene rare-vs-common PTC split as an internal control. Darker shading indicates the FDR-passing fraction, lighter shading the nominally-significant-only fraction. (g) Inverse-variance–weighted meta-analysis of per-gene log OR of NMD-triggering to NMD-evading somatic PTCs from TCGA. For each category two estimates are shown: rule-based NMD classification (filled circles) and rule-based with NMDetective-AI concordance filter (open squares). Whiskers show 95% confidence intervals. The gray band indicates the 95% CI of the “Other Genes” background. Fixed-effects (FE) or random-effects (RE; DerSimonian-Laird) pooling is applied depending on heterogeneity (RE when I² > 50%). (h) Per-gene log odds ratio of NMD-triggering to NMD-evading somatic PTCs (normalized by synonymous burden) for individual tumor suppressor genes (*MUC4* not shown). Points show Wald log OR with 95% CI for genes with nominal *p* < 0.05. FDR-significant genes (Benjamini-Hochberg q < 0.05) are highlighted. Colors indicate direction of selection. Dashed horizontal line at y = 0 represents no enrichment.

Comparing the rare gnomAD variants versus a set of common gnomAD variants (AF ≥0.1% in ≥1 subpopulation; n = 2,649) revealed differential selection on NMD categories (Fig. 8d). Common PTCs showed a depletion of NMD-triggering variants (42.7% vs 54.0% in rare; Fisher’s exact test *p* = 1.38e-30) and corresponding enrichment of NMD-evading categories, particularly last exon variants (36.2% vs 24.1%) and start-proximal variants (9.4% vs 8.2%). This indicates that NMD-triggering PTCs are overall under stronger purifying selection.

### Differential NMD activity on variants in disease genes can inform clinical decision making

To assess NMD variant composition in individual disease genes, we analyzed ClinVar disease genes with ≥10 unique rare PTCs in gnomAD, using two complementary enrichment tests (based on Wald log OR statistical tests, Fig. 8e-f; see Methods). Test 1 compares rare PTC allele counts in NMD-triggering versus NMD-evading regions, normalized by synonymous and missense allele counts in the same regions; Test 2 compares NMD-triggering versus NMD-evading region, contrasting rarer versus more common PTCs within each gene. Of 714 tested genes, 260 (36%) showed no nominal significance in either test (Fig. 8f). Among NMD evading-enriched genes (i.e. those where NMD, overall, associates with stronger negative selection and we thus infer NMD would aggravate disease phenotype), 167 passed test 1 FDR < 1%; as a validation analysis, of these, 56 genes also passed test 2 FDR < 25%. Among the converse case of NMD triggering-enriched genes (i.e. those where NMD associates with weaker negative selection, and thus likely alleviates phenotypes), 129 passed test 1 FDR and 38 genes also passed the test 2 validation thresholds. Considering the genes where both tests were significant, among the genes with the largest effect sizes of NMD-triggering depleted genes in broadly healthy populations (Fig. 8e), here we highlight *CTC1* telomere maintenance gene, causing the pleiotropic Coats plus syndrome (log base e OR test 1 = −7.36, FDR corrected combined *p* = 1.4e-98), the *WRN* helicase, causing Werner syndrome adult progeria (log OR test 1 = −5.70, FDR corrected combined *p* = 4.3e-143) and *CNGB1* retinitis pigmentosa disease gene (log OR test 1 = −4.64, FDR corrected combined *p* = 1.5e-89). There were also large effect size genes among NMD triggering-enriched genes (Fig. 8e). Both directions of NMD imbalance are broadly represented among known disease genes, with a quantitatively higher prevalence of the putatively NMD-aggravating genes, suggesting that as far as genetic disease goes NMD should not be considered an overall protective pathway. The rigorous, two-stage testing setup allowed us to identify individual genes with higher confidence in the NMD effect direction (full list in Table S4), opening avenues for prioritizing NMD inhibition and stop codon readthrough therapies in the corresponding diseases. We noted strong agreement between log OR scores from test 1 and test 2 for genes passing the FDR criteria (R=0.89), and more moderate agreement across all tested genes (R=0.39; Fig. S16), likely in part due to statistical power.

Next, we investigated NMD effects on positive selection patterns in somatic mutations, to systematically assess the role of NMD in cancer genome evolution. We performed a per-gene analysis of somatic PTCs from TCGA whole-exome sequencing, testing enrichment in NMD-triggering versus NMD-evading regions using variants with NMDetective-AI support, and with synonymous variants in the same regions as a local mutation rate control (see Methods; Fig. 8g-h). Per-gene estimates were pooled within gene categories using inverse-variance–weighted meta-analysis (Fig. 8g; see Methods). Tumor suppressor genes (n = 84 genes with sufficient data) showed a pooled log OR of 0.52 (95% CI: 0.34–0.70; random-effects model; *p* = 1.3e-8), significantly exceeding the “Other genes” background set (log OR = 0.23; *p* diff = 1.4e-3), signaling positive selection for loss-of-function through NMD-mediated transcript degradation. Essential genes (n = 227) showed the opposite trend,, with a pooled log OR of −0.07 (95% CI: −0.16 to 0.02; *p* = 0.11 vs zero), significantly below the “Other genes” background (*p* diff = 1.9e-10), supporting negative selection against NMD-triggering PTCs that would eliminate essential gene function in tumor cells^6^. Oncogenes (n = 22) showed no significant signal (log OR = 0.06, 95% CI: −0.21 to 0.33; *p* = 0.67). Using rule-based NMD classification alone yielded qualitatively consistent but attenuated results (TSG log OR = 0.39, *p* = 3.5e-7), confirming that relying on NMDetective-AI enriches for bona fide NMD signal.

At the individual gene level, we computed per-gene log ORs for 178 tumor suppressor genes with sufficient somatic PTCs and synonymous variants (Fig. 8h). Of these, 29 showed significant NMD category imbalance after FDR correction (Benjamini-Hochberg q < 0.05): 24 enriched for NMD-triggering PTCs and 5 enriched for NMD-evading PTCs. The largest effect sizes among triggering-enriched genes shown in Fig. 8h were *SDHA* (log OR = 2.59), *RB1* (log OR = 2.59), *MAP2K4* (log OR = 2.52), and *KDM5C,* all well-established tumor suppressors. For these and other similarly enriched genes (Fig 8h), we infer that NMD-mediated loss of function is a major pathogenic mechanism; the implication is that tumors bearing driver mutations in these genes would be prioritized for applying NMD inhibitors and PTC readthrough agents. Among the rarer cases of NMD evading-enriched significant cancer driver genes (q < 0.05), the largest effect sizes were observed in tumor suppressors WRN (log OR = −2.32), *HLA-B* (−1.62), *PTCH1* (−1.61), and in the common tumor suppressor *NF1*, suggesting that truncated protein products in these genes may act via dominant-negative mechanisms, or, more speculatively, other gain-of-function mechanisms.

## Discussion

Building on our NMDetective framework^6,22^, which applied genome-wide NMD rules to interpret genetic disease and cancer driver mutations (as well as implications of NMD to CRISPR editing and cancer immunotherapy), here we move from rule-based annotations toward quantitative prediction of variant-specific NMD efficiency. Large-scale genomics studies have already shown that the canonical positional rules capture real signal but leave substantial unexplained variability^20,24^, which we estimated to be roughly ∼1/3^22^. Our results suggest that the limitation of prior frameworks is not that the rules are inaccurate or grossly incomplete, but that their outcomes are treated as categorical and that their variation across genes is not modelled. In our genome-wide analyses and DMS experimental data, the classical 50-nt penultimate exon rule is better represented as a graded transition with a mean inflection point close to the classical boundary and showing between-gene heterogeneity. Similarly, for the long-exon rule previous studies inferred a >∼400-nt effect from genomic, observational data^6,22^ and implemented it as an annotation rule^1,20,23,32^; our mutagenesis data provide direct experimental support and show that exon length and PTC position interact quantitatively, with progressive evasion in long exons and increasingly flat, globally weak NMD in very long exons. A similar distinction emerges for the start-proximal region NMD escape. Early studies of the rule on individual examples of PTCs in disease genes^10–13^, as well as transcriptome-scale analyses^6^ supported a broad 5′ escape zone. A recent saturation-editing study on the human *LMNA* gene^55^ suggested that gene-specific boundaries on 5’ evasion could be sharp, and averaging over many genes would create a ramp in the overall rule. Our multi-gene, comprehensive DMS results are consistent with that interpretation and extend it in two ways: first, by showing substantial heterogeneity of start-proximal NMD escape profiles across many genes, and second, by directly perturbing translation reinitiation capacity. Downstream AUGs increased RNA abundance by about twofold in our reporter, with effects depending on intercistronic distance, uORF length and Kozak context, and with direct protein evidence for downstream reinitiation.

Our computational model based on neural networks, NMDetective-AI, trained and tested on large scale human genomic and transcriptomic datasets, broadly recapitulates these refined NMD rules in the penultimate exon 50 nt region, in long exons and also in the start-proximal region. For comparison, recent computational models of NMD activity, including our previous NMDetective-A/-B^22,32,56^ rely on fixed NMD rules based on PTC and transcript features, or a regression model fit to the same features. NMDetective-AI instead does not rely on any preconceived features, rather it learns them in a de novo, unbiased manner, starting from the mRNA sequence and annotations of splice site positions, based on the versatile Orthrus RNA foundation model^28^. The contemporary NMDEP method^57^ also relies on Orthrus albeit in a different way, using neural net-derived sequence representations (embeddings) of mRNA, notably concluding they are well predictive only when complemented with curated features such as evolutionary conservation, ribosome loading, and known NMD rules. By contrast, our results indicate that end-to-end fine-tuning of the Orthrus RNA language model on the full transcript sequence and structure, deriving NMDetective-AI, can extract additional quantitative signal beyond fixed rules, supporting nucleotide-resolution, gene-aware prediction. Our study shows that a sequence model, when coupled to large-scale experimental validation, can capture the local and gene-specific structure of NMD efficiency that hand-crafted features necessarily compress. Immediate sequence context is likely part of that signal, consistent with both our own DMS scores residual analysis and two recent experimental studies^49,55^. These involve neighboring amino acids and also local readthrough-permissive sequence context as NMD modifiers superimposed on positional rules. While their effect is overall subtle, in individual examples of PTC the local context may play a substantial role: in our DMS experimental readouts, neighboring PTC may have variable NMD efficiency in a way that reproduces across the different PTC codons, suggesting further local sequence variations whose effect on NMD and mechanisms thereof remain to be elucidated going forward.

While the neural net model is complex and a “black-box”, the *in silico* mutagenesis performed on NMDetective-AI can be used to generate dense NMD data points, appropriate for deriving simple yet accurate models of local NMD efficacy ramps; we demonstrate this via fitting 4-parameter logistic curves for individual segmental rules. These interpretable representations of NMDetective-AI may further mechanistic studies, and provide transparency in predictions, important for genetic medicine applications.

More broadly, our study bridges two long-standing views of NMD: a mechanistic framework built from gene-specific reporter studies, and a statistical framework derived from population and cancer genomics. By combining both with mRNA sequence-based modelling to generate the NMDetective-AI, highly accurate NMD predictive model, we show that the classical NMD rules remain biologically meaningful but are only approximations to a richer transcript-level landscape. This has practical consequences for variant interpretation and for therapeutic stratification, because whether NMD protects from or contributes to disease cannot be inferred from the logic of the NMD pathway alone, but needs to be resolved at the level of individual genes and variants.

## Methods

### Deep mutational scans - library design and cloning

#### 50nt rule library

Oligos were designed to encode all stop codons and tryptophan (UAG, UAA, UGA and UGG) substitutions for the 3’-most 76 amino acid positions of the penultimate exon in BRCA1 and ATP7A minigenes, expressed from a mammalian expression vector (pIT078-BRCA1 and pIT078-ATP7A, Table S5. UGG variants were used as 0% NMD control. BRCA1 mutated exon is exon 10, and it’s flanked by a VDJ fragment upstream (to prevent the start proximal NMD evasion) and BRCA1 intron 10 and exon 11 downstream (to provide an EJC to trigger NMD). ATP7A minigene contains exons 2-3, intron 3, exon 4, intron 4 and exons 5-6, and the library covers the 3’ end of exon 4. Oligos were ordered as an oligopool to Twist Biosciences containing the variable part (library) and gene-specific flanking constant sequences for PCR amplification. The oligopool was PCR-amplified for 14 cycles and cloned into pIT078-BRCA1 and pIT078-ATP7A minigenes using Gibson assembly. The library was sequenced with Illumina MiSeq 300 paired-end reads.

#### Long exon rule library

We generated nine 5’ truncated versions of BRCA1 exon 10 using Gibson Assembly. Then, we used nicking mutagenesis^45^ to mutate all VYY (where V=A,C or G, and Y=C or T) codons (n=83) to all three stop codons and UGG, yielding a total of 950 variants. UGG variants were used as 0% NMD control. Oligos used for nicking mutagenesis were ordered to Integrated DNA Technologies (IDT). The library was digested out from the nicking mutagenesis plasmid using XhoI and pspOMI restriction enzymes and cloned into the same mammalian expression plasmid pIT078. As described above for the 50nts rule experiment, pIT078 drives the expression of a minigene with a VDJ fragment upstream of the library to prevent the start proximal NMD evasion, as well as BRCA1 intron 10 and exon 11 downstream, to provide an EJC binding site to trigger NMD. The library was sequenced with a MinION (ONT, R9.4.1 flow cells).

#### Start proximal rule libraries

For the BRCA1 library, oligos were designed as explained in Main text and ordered as an oligopool to Twist Biosciences containing the variable part (library) flanked by constant sequences for PCR amplification. The oligopool was PCR-amplified for 14 cycles and cloned into pIT075 plasmid using Gibson assembly. pIT075 harbors exons BRCA1 2-11 and introns 5, 9 and 10 (Fig. 6a). The library covers the 5’ most 83 BRCA1 residues.

We designed a second library by generating all UAG variants across the 5’ -most 83 residues (plus the non-PTC WT) for the 139 genes with highest number of pathogenic nonsense mutations in the human population across genetic diseases and cancer (ClinVar, MSK-Impact, TCGA). The gene selection process was as follows: genetic disease and cancer germline variants were retrieved from the ClinVar database, where all pathogenic nonsense variants whose review status was two or more stars were included (n = 3498). Somatic cancer variants were obtained from MSK-IMPACT and TCGA databases with at least two entries in either dataset (n = 2,372). We then clustered variants by gene and selected the 139 genes with the highest number of nonsense mutations. The library was cloned into the *BRCA1* minigene (Fig. 6a). The library was sequenced with Illumina MiSeq 300 paired-end reads.

#### Library transformation

Libraries were transformed using Neb10 Electrocompetent bacteria and grown in 100mL overnight culture. Library complexity was estimated by plating a small amount of the transformation reaction and extrapolating the total number of transformants. Individual clones were Sanger sequenced to confirm the expected structure and diversity.

### Deep mutational scans - sequencing data generation and analysis

#### Cell culture and transfection

HeLa cells were maintained in DMEM supplemented with 10% FBS without antibiotics and grown at 37°C at 5% CO2. 3e^-05 cells seeded in 6-well plates were transfected with 1 μl Lipofectamine 3000, 1 μl P3000 reagent and 150ng of library DNA. 48h later, RNA was harvested using RNeasy (Qiagen). During these 48h the frequency of variants change according to their NMD sensitivity. Note that before the library experiment, all NMD reporters used in the study were tested with 2-3 PTCs to ensure they were sensitive to NMD.

#### Library preparation for Illumina Sequencing

RNA was extracted from HeLa cells using the RNeasy kit (Qiagen), eluted in 30ul of MilliQ, and treated with TurboDNase (Thermo Fisher) to degrade residual genomic DNA together with RNAseIN to prevent the action of RNAses. To avoid freeze/thaw cycles, we performed the cDNA synthesis right after. We used 1.5ug of RNA together with a gene-specific primer (oIT203, Table S5) that anneals at the library-specific 3’UTR to convert the library transcripts to cDNA. The cDNA samples are referred to as ‘outputs’, since they belong to the library after NMD selection. In parallel, we prepared DNA aliquots from the library MIDIpreps for sequencing as inputs.

We quantified the DNA concentration (plasmid DNA or cDNA, for inputs and outputs; respectively) in triplicate by real-time quantitative PCR, using primers that had homology to the origin of replication region of the minigene’s plasmids backbone (input) or to *BRCA1*/*ATP7A* ORF regions (output).

A 2-step PCR using high fidelity Q5 Hot Start High-Fidelity DNA Polymerase (NEB) was used to amplify the input and output libraries for sequencing. In PCR1, ∼50 million molecules were amplified for 15 cycles using frame-shifted primers (to improve sequencing performance) with overhang homology to Illumina sequencing adapters (oIT_ILL_01/03/07-12, Table S5). These primers were specific to each library, and PCR settings were optimized for each primer pair. Then, PCRs were purified using the MinElute PCR Purification Kit (QIAGEN) according to the manufacturer’s protocol. DNA was eluted in Milli-Q water to a volume of 20ul and quantified with Nanodrop. Next, 30ng of PCR1 product was used as template for PCR2. Here, the remaining parts of the Illumina adapters were added to the library amplicon. The forward primer (oIT_GJJ_1J) was the same for all samples, while the reverse primer (oIT_GJJ_2J) differed by the barcode index. 8 cycles of PCR2s were run at 62°C of annealing temperature and 25 seconds of extension time. An aliquot of each PCR2 was run on a 2% agarose gel to be quantified. After quantification, samples with different Illumina indexes that were sequenced together in the same flow-cell were pooled in an equimolar ratio, run on a gel and purified using the QIAEX II Gel Extraction Kit. The purified amplicon library pools were subjected to Illumina 300bp paired-end MiSeq sequencing at the CRG Genomics Core Facility.

#### Library preparation for ONT Sequencing

Due to the amplicon length, the long exon rule libraries were sequenced with ONT, which required a different library preparation than Illumina. Variants were grouped by exon length, and the experiment was performed independently across groups to prevent length-related transfection, purification and PCR biases. Similar to the Illumina-sequenced libraries, RNeasy kit (Qiagen) was used to extract RNA and TurboDNase (Thermo Fisher) was used to degrade residual genomic DNA. 1.5ug of RNA together with a gene-specific primer that anneals at the library-specific 3’UTR were used to convert the library transcripts to cDNA.

Primers targeting the constant sequences flanking the libraries were used to PCR-amplify the library transcripts from the plasmid (input, oIT272-277, Table S5) and the cDNA pools (output, oIT330-335, Table S5), using an annealing temperature of 67°C, 4ul of cDNA as template and the Q5 Hot Start High-Fidelity DNA Polymerase (NEB). The extension time (30’’ to 6’) and number of cycles (21-24 cycles) were adjusted for each exon length. Primers were barcoded to allow demultiplexing of replicates. Samples were purified using the QIAEX II Gel Extraction Kit and quantified by qPCR. To sequence variants of different lengths in the same run and get a homogenous coverage, we pipetted the same number of molecules of each length bin into the flow cell. To further control for the amplicon length sequencing bias, we sequenced the libraries in two runs: a) exon lengths 125bps-1500bps and b) exon lengths 2500-3426bps. For library prep we used the Ligation Sequencing Kit (SQK_LSK109) according to the manufacturer’s protocol, which repairs the DNA and attaches the sequencing adapters. Next, MinION R9.4.1 flow cells were primed and sequencing lasted for 24h, until getting ∼5M reads. Inputs and outputs were sequenced separately (total of 4 ONT runs and 2 flow cells, since they were washed in between runs and reused).

#### Sequencing data processing for Illumina Sequencing

FastQ files from paired-end sequencing of all experiments were processed with DiMSum (v.1.3) to obtain the read counts and NMD efficiency for each variant^46^. DimSum applies stringent quality filters to discard low quality reads, reads with sequencing errors, etc. to ensure that only high-quality reads are used for downstream analysis. Experiments were performed in biological triplicates (except for the ATP7A 50-55nts rule library, which was performed in duplicates), and we retained variants with an average, across all inputs and outputs, of >=10 reads for downstream analyses. Increasing the >=10 reads filter did not improve inter-replicate correlation. Median coverage in all libraries is >130 reads per variant. Also, note that for some libraries, such as the 50-55 nts and the SPR in 122 genes all variants are sequenced with >40 reads.

In DMS experiments, the enrichment score (ES) for each variant i in replicate r is usually defined as the ratio between its frequency before and after selection, where N^input^ and N^output^ are the corresponding numbers of sequenced reads in the input and output samples, respectively:

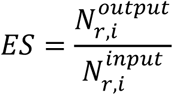

DiMSum calculates RNA levels scores for each variant i in replicate r as the natural logarithm of the enrichment score normalized to the wild-type variant (wt):

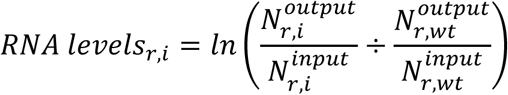

Errors are estimated by fitting a modified Poisson (count-based) model to the data, with replicate-specific multiplicative and additive terms that are common to all variants. Final DiMSum RNA levels scores and error estimates for all variants are obtained by merging between replicates using weighted averaging, whereby fitness of low-error replicates weigh more than high-error replicates.

#### Sequencing data processing for ONT Sequencing

Fast5 files were base called with Guppy 5.0.5 using the ‘hac’ mode and only reads that passed the quality filters were retained. Porechop was used to demultiplex the replicates based on the replicate-specific barcodes (https://github.com/rrwick/Porechop). Then, reads were mapped to each of the nine exon lengths using the SeqKit package and aligned to the reference WT sequences using minimap2. Sam2tsv (Jvarkit) was used to convert the SAM alignments to a TAB delimited file, and an in-house script was used to quantify the number of full-length reads for each variant. Same as for Illumina libraries, we calculated RNA levels scores for each variant *i* in replicate *r* as the natural logarithm of the enrichment score normalized to the wild-type variant. Experiments were performed in biological triplicates, and we retained variants with >=10 reads.

#### qPCR

We used qPCR to validate the DMS results for the start proximal library. Cells were transfected with 200ng of the BRCA1 validation colonies (n=22) and 200ng of a transfection control plasmid encoding for the VDJ gene. 48h later, RNA was extracted from HeLa cells using the RNeasy kit (Qiagen), eluted in 30ul of MilliQ, and treated with TurboDNase (Thermo Fisher) to degrade residual genomic DNA together with RNAseIN to prevent the action of RNAses. To avoid freeze/thaw cycles, we performed the cDNA synthesis right after. We used 1.5ug of RNA together with a gene-specific primer that anneals at the library-specific 3’UTR to convert the library transcripts to cDNA. qPCR was run with primers amplifying the VDJ control gene and primers amplifying the BRCA1 gene. VDJ was used as a transfection control to normalize the BRCA1 signal.

#### Western Blotting

Cells were transfected with 200 ng of either BRCA1-WT or BRCA1 start-proximal library, expressing an alpha-tag at the 3’end of the transcript. 48h later, cells were trypsinized, washed with PBS (x3), incubated with lysis buffer (+ protease&phosphatase inhibitors) and sonicated for 5’. 20ul of sample and Laemli loading buffer (3:1) were run on a gel and transferred to a nitrocellulose membrane using iBlot. After 1h blocking incubation with 5% Milk in TBS-T we incubated overnight with anti-actin (Thermo Fisher) and anti-alpha tag (FluoTag®-X2 anti-ALFA, NanoTag) antibodies. The anti-alpha tag membranes were imaged on the Odyssey CLx Imager. The actin membrane was incubated for 1h with a secondary antibody and bands were imaged on an iBright CL1500 Imaging System.

### Processing of genomic and transcriptomic data

#### Source Datasets

We obtained premature termination codons (PTC) from nonsense and indel variants with allele-specific expression (ASE) measurements from three sources: somatic mutations from The Cancer Genome Atlas (TCGA, 8,786 tumor samples, 31 cancer types), germline variants from TCGA (9,752 individuals), and germline variants from the Genotype-Tissue Expression (GTEx, 838 individuals, 15,247 healthy samples, 56 tissues) project^42^. For this, we downloaded matched tumor and normal WES *bam* files, along with tumor RNA-seq *fastq* files, from the TCGA consortium via the Genomic Data Commons (GDC) Data Portal (https://portal.gdc.cancer.gov). From GTEx project, we directly obtained WES VCF and ASE counts files for the 838 individuals, accessed through dbGaP (via AnVIL, https://anvilproject.org/), v8 (dated 05–06-2017).

#### RNA-seq alignment

We used the same methodologies as in our recent paper^1^, here summarized for convenience. We processed TCGA RNA-seq data by aligning the reads to the human genome *GRCh38.d1.vd1*, utilizing *STAR v2.5.3a*^58^, in adherence to GTEx guidelines (https://github.com/broadinstitute/gtex-pipeline/tree/master/rnaseq). Only RNA-seq reads achieving a mapping quality score of at least 255 were kept. We used WASP^59^ to correct for allele-mapping bias and filter out duplicate reads.

#### Variant calling, annotation and ASE quantification

Variant calling for TCGA whole-exome sequencing (WES) and separately for RNA-seq data was performed using Strelka2 v2.9.10^60^. Somatic variants were identified by comparing paired tumor and normal BAM files, while germline variants and RNA-seq ASE counts were derived solely from normal BAM files, with RNA-seq variant calls forced to match WES-derived calls. SNVs and indels were called separately, merged, and processed through an in-house script to quantify ASE (available at: https://github.com/gpalou4/iNMDeff). All resulting VCFs were annotated using ANNOVAR^61^ (2020–06-07 version, GENCODE v26, ENSEMBL v88), incorporating gnomAD^62^ for population minor allele frequencies (MAF). For GTEx data, pre-computed WES VCFs and ASE counts were downloaded directly, converted from GRCh37 to GRCh38 coordinates via liftover (excluding regions with poor alignment^63^), and annotated using the identical ANNOVAR pipeline.

#### Quantifying raw PTC NMD efficiency

Raw NMD efficiency was quantified using allele-specific expression (ASE) from RNA-seq data. For each heterozygous PTC variant, the ASE-based metric represents the negative log_2_ of the ratio in mRNA expression of the mutant (PTC) allele relative to the reference allele. In this raw score, 0 indicates no detectable NMD activity, a value of 1 corresponds to a 50% reduction in mRNA level, while a complete heterozygous mRNA degradation approaches infinity. Raw NMD scores were further processed before training, see later.

#### Filtering and processing of variants - shared filters

Somatic and germline mutations from the TCGA dataset, alongside germline variants from the GTEx project, were processed using a unified filtering framework (Supp. Fig. 1) with cohort-specific adaptations. Across all three datasets, samples exhibiting extreme mutational burden ratios (a nonsense-to-indel or deletion-to-insertion ratio > 10 or < 0.1) were excluded. Transcripts were assigned a priority score based on ENSEMBL Transcript Support Level (TSL) and MANE main isoform status (1: TSL = 1 and MANE; 2: TSL = 1, non-MANE; 3: TSL > 1, MANE; 4: all others), retaining only transcripts with a priority score ≤ 3. To isolate true NMD effects, PTCs with conflicting classifications across multiple transcripts due to alternative UTR structures were removed. For variants overlapping multiple transcripts, a single representative transcript was selected based on the previous priority hierarchy (TSL/MANE score = 1). Similarly, we excluded PTCs that manifested as missense, synonymous, or non- coding UTR mutations in alternative transcripts due to differing reading frames. Frameshift indels lacking a predicted downstream PTC were discarded. To prevent confounding effects from multiple truncating mutations, variants were excluded if the patient harbored an additional germline loss-of-function variant (nonsense, frameshift indel, or splice site variant) within the same gene. We also removed variants occurring in single-exon genes – unless they contained a 3’ UTR splice site – as these typically evade canonical NMD. Only heterozygous variants were retained to ensure clear allele-specific analysis. Finally, variants resulting in undefined (NA) or infinite values during NMD efficiency calculations due to insufficient RNA-seq coverage were removed.

#### Filtering and processing of variants - dataset specific filters

Depending on the dataset, specific filters were applied or omitted to account for variant origin and tissue context. For both TCGA datasets (somatic and germline), PTCs occurring within multi-nucleotide variants (MNVs) were excluded to prevent “MNV rescue” where an adjacent mutation neutralizes the stop-gain effect. Furthermore, PTCs overlapping somatic focal copy number alterations (absolute GISTIC score > 0.1) were excluded to prevent copy number artifacts from confounding expression calculations. Both the MNV and CNA filters were omitted for the GTEx dataset. To guarantee high-confidence calls for tumor-specific mutations, the somatic TCGA pipeline filtered out variants with a DNA VAF < 0.2. This DNA VAF filter was omitted for both the TCGA and GTEx germline datasets. Expression-based filtering was applied across all datasets to ensure robust baseline expression for ASE analysis. For the germline datasets, transcripts with low expression (median mRNA expression < 5 TPM) or high variance (coefficient of variation [CV] > 0.5) were removed at this stage. Because the Somatic TCGA cohort served as the training dataset, these expression filters were applied in a subsequent processing step utilizing less stringent thresholds (TPM < 1 and CV > 1). Furthermore, common PTCs (MAF > 0.1%), singletons (MAF = 0.0%) and PTCs within 3 nucleotides of splice sites were excluded.

#### Additional preprocessing before modeling

We performed further filtering and preprocessing steps on the initial NMD dataset, before model training. For germline datasets, variants overlapping with somatic TCGA PTCs were removed using both genomic coordinates and transcript sequences to ensure test set independence. Additional filtering based on transcript expression and variation were used for PTCs on validation chromosomes (TPM ≥5.0, CV ≤0.5).

To isolate sequence-dependent NMD effects from biological and technical confounders, we performed linear regression correction separately for stopgain and frameshift variants. Predictor variables consisted of the first four principal components (PC1–PC4) derived from tissue-specific RNA-seq gene expression data, the first principal component (PC1) of mRNA half-lives from multiple experimental studies^64^, and expression metrics (median TPM, CV). Missing RNA half-life values were imputed with the dataset median. Frameshift-induced PTCs were corrected using a two-stage adjustment to match stopgain variant levels.

Variants sharing identical characteristics (gene, transcript, position, sequences, stop codons, rule labels) i.e. recurrent observations of the same PTC event across samples were aggregated using median NMD efficiency. Extreme values (|NMDeff| > 4.0) and transcripts exceeding 20,000 nucleotides were filtered from training data. NMD efficiency values were normalized using a linear transformation where the center point was the midpoint between mean NMD-triggering and mean last-exon variant efficiency, and the scale was their difference, in essence setting the value of full NMD triggering to 0.5, and full NMD evasion to -0.5. Normalized values were clipped to [-2, 2]. This normalization was performed independently for each dataset.

For each variant, sequences were encoded as 6-track one-hot representations: four nucleotide tracks (A, C, G, T), one CDS track marking codon boundaries, and one splice site track indicating exon-exon junctions. All preprocessing configurations and statistics were saved for reproducibility. The 3’ and 5’ UTRs for each transcript were obtained using the Genome kit library with Gencode v26.

#### Harmonizing DMS datasets with large-scale genomic data

DMS datasets required specialized preprocessing to align experimental fitness measurements with TCGA NMD efficiency scales.

#### Penultimate exon dataset

Variants with DimSum σ > 0.15 or RNA abundance levels < −2 were removed. Fitness values were mean-centred and inverted for visualization purposes. Normalization was performed to the mean and standard deviation of NMDetective-AI predictions in the same regions of the same genes for visualization (Fig. 4b-c). Two genes were included: BRCA1 (ENST00000357654.7) and ATP7A (ENST00000341514.10).

#### Long exon dataset

Measurements were first mean-centred, and values outside [−1, 1] were excluded. Fitness was inverted (NMDeff = −fitness) with no further normalization for visualizations only. Target: BRCA1 (transcript ENST00000357654.7, exon 10).

#### Start-proximal exon dataset

Variants with DimSum σ > 1, and variants with |measurements| > 3 were dropped. Genes where >50% of variants had higher fitness than the wild-type measurement were discarded as likely confounded, as were genes with fewer than 50 remaining variants. RNA level was inverted to yield NMDeff (NMDeff = −fitness). Normalization proceeded in two steps: first, for each gene, the maximum rolling median across the top-15 3′-most variants (in three groups of 5 variants) was shifted to +0.5 (the expected 3′ NMD-evading anchor); second, a global linear scaling was applied, anchored at +0.5 on the 3′ end, to match a target 5′ median of −0.5. Variants mapping to last-exon positions were excluded.

### Model development and training

#### Model architecture

NMDetective-AI is a deep learning model that combines the pretrained genomic foundation model Orthrus with a regression head for sequence-based prediction of NMD efficiency. The model is trained end-to-end, with no frozen components. The architecture consists of: Encoder: We used Orthrus^28^ through HuggingFace (https://huggingface.co/antichronology/orthrus), a Mamba-based mature RNA foundation model with approximately 10 million parameters, pretrained on over 45 million mature RNA transcripts from mammalian organisms. Orthrus encodes full-length transcript sequences into 512-dimensional embedding vectors that capture functional and evolutionary relationships across sequences. The model processes input sequences in a 6-track one-hot encoding format: four nucleotide tracks (A, C, G, T/U), one coding sequence (CDS) track marking exon boundaries, and one splice site track indicating exon-exon junctions. We generate representations using mean pooling over the sequence length. Regression Head: The Orthrus embeddings are passed through a deep neural network consisting of two fully connected layers (256 and 64 neurons) with GELU activation functions, layer normalization after the first layer, and dropout regularization. The final layer outputs a single scalar value representing predicted NMD efficiency. Weight initialization for the regression head used Xavier uniform initialization for linear layers, with biases initialized to zero. All models were implemented in PyTorch^65^.

#### Hyperparameter optimization

Hyperparameter optimization was performed using Bayesian optimization via Weights & Biases (W&B)^66^ sweep with 42 instances. The hyperparameter search space included:

- Learning rate: [0.00001, 0.00005, 0.0001, 0.001]
- Weight decay (head): [0, 1e-5, 1e-3]
- Weight decay (encoder): [0, 1e-5, 1e-3]
- Learning rate decay gamma: [0.9, 0.95, 0.99]
- Gradient accumulation steps: [8, 64, 256]
- Regression head architecture: [[256, 64], [512, 256, 128, 64], [1024, 512, 256, 128, 64]]
- Dropout: [0.0, 0.1, 0.2]
- Gradient clipping norm: [1.0, 2.0, 3.0]
- Warmup steps: [50, 100, 10000]
- Encoder version: [standard Orthrus, MLM-pretrained Orthrus]
- Activation function: [ReLU, GELU]
- Layer normalization: [True, False]
- Loss function: [MSE, Huber]
- Huber delta: [0.5, 1.0, 2.0]

The optimization objective was to minimize validation loss. The best-performing configuration was selected based on validation set performance. Logs for the sweep can be viewed at https://wandb.ai/irb-gds/NMD/sweeps/xzhwjczx.

#### Training strategy

The final NMDetective-AI model was trained in the following manner.

Optimization: Models were trained using AdamW^67^ optimizer with separate weight decay parameters for the encoder (0.001) and regression head (0.0). Gradient clipping was applied with a maximum norm of 3.0.

Training Configuration: Training used a batch size of 1 with gradient accumulation over 256 steps, yielding an effective batch size of 256. The model was trained for a maximum of 300,000 steps with evaluation performed every 1,024 steps. Huber loss (delta=0.5) was used as the training objective. Early stopping was implemented with patience of 50 evaluation steps without improvement in validation loss.

In addition, we evaluated three training strategies to assess the contribution of pretrained Orthrus representations: (1) NMDetective-AI: full fine-tuning of the entire model including the encoder, (2) linear probing with frozen encoder weights training only the regression head, and (3) random initialization of the encoder without pretraining. When training models (2) and (3) the same configurations were used. Logs for trained models can be inspected at https://wandb.ai/irb-gds/NMD.

### Other statistical analysis

#### In silico PTC scan experiments

To systematically evaluate NMDetective-AI predictions across transcript sequences, we performed in silico saturating mutagenesis experiments. For selected transcripts, we computationally introduced premature termination codons at positions of interest within the coding sequence by replacing codons with stop codons (UAG, UGA, or UAA). For each variant, the full-length mutated transcript sequence was encoded in 6-track format and passed through NMDetective-AI to generate NMD efficiency predictions. This approach allowed comprehensive profiling of positional effects on NMD across parts of or entire transcripts, enabling comparison with experimentally measured deep mutational scanning data and validation of model predictions across diverse sequence contexts.

#### Reimplementing NMDetective-A and NMDetective-B

To compare NMDetective-AI with previous approaches, we reimplemented NMDetective-A and NMDetective-B using our normalized NMD efficiency labels. NMDetective-A is a Random Forest regressor (100,000 trees, max_features=1) trained on seven hand-crafted features: InLastExon (boolean), 50ntToLastEJ (boolean), DistanceToStart (capped at 1000 nt), ExonLength, PTC_EJC_dist (distance to downstream exon-exon junction), DistanceToWTStop, and RNAHalfLife. NMDetective-B is a decision tree with fixed branching structure based on four features (InLastExon, DistanceToStart <150 nt, ExonLength >407 nt, 50ntToLastEJ), where leaf node values are computed as mean NMD efficiency from training data for variants matching each decision path. We also evaluated NMDetective-B with original fixed leaf values from the published model. All features were pre-computed during the preprocessing pipeline described above.

#### Measuring inter-replicate correlation

To assess measurement reproducibility, we performed split-half reliability analysis on PTCs with multiple biological replicates (≥3 independent samples with the same PTC). For each PTC, replicates were randomly divided into two groups, and mean NMD efficiency was calculated per group. Spearman correlation was computed across all PTCs between the two group means. This random splitting was repeated 100 times to obtain robust estimates of reproducibility. The analysis was performed separately for NMD-triggering variants (meeting none of the escape rules) and NMD-evading variants (meeting at least one escape rule). To compare measurement consistency across cohorts, we calculated correlations between ASE measurements for the same PTCs appearing in different datasets (somatic TCGA vs germline TCGA, somatic TCGA vs GTEx).

#### Fitting LOESS, and four-parameter logistic curves

LOESS (locally weighted scatterplot smoothing) was used to fit smooth curves to positional NMD efficiency data. For DMS start-proximal analysis, LOESS curves (frac = 0.3) were fitted per gene to NMD efficiency as a function of PTC position, then linearly interpolated at 83 uniformly spaced positions (1–83 codons from the start, corresponding to CDS positions 3–249 nt), creating a gene × position matrix used as input for PCA and hierarchical clustering. For the penultimate exon normalization (DMS PE dataset), LOESS was applied with frac = 0.4 to align DMS and TCGA PTC distributions.

Four-parameter (Log-)logistic (4PL) curves were fitted to characterize the relationships in the penultimate exon data (genome-wide predictions). The 4PL function is defined as:

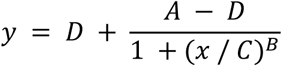

where A is the minimum asymptote, D is the maximum asymptote, C is the inflection point, and B is the slope factor. x was normalized to [0, 1] for numerical stability. Curve fitting used ‘scipy.optimize.curve_fit’ with Levenberg–Marquardt, starting from data-derived initial guesses, with a maximum of 20,000 function evaluations.

Boltzmann sigmoid or logistic curves of the form:

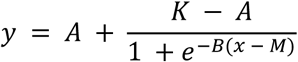

where A is the lower asymptote, K is the upper asymptote, B is the growth rate, and M is the midpoint (inflection point), were fitted to DMS start-proximal data on a per-gene basis. x was scaled to [0, 1] prior to fitting. Bounded least-squares optimization with a soft-L1 loss function (f_scale = 0.5, max_nfev = 40,000) was used via ‘scipy.optimize.least_squares’. Orientation of the sigmoid (increasing vs. decreasing) was detected from the data and used to initialize parameters. Genes for which fitting failed were excluded from downstream sigmoid-based analyses. Goodness of fit was assessed by R².

For the long exon dataset, multiple curve models were compared per exon: linear, 4-parameter logistic, and polynomials of degree 2–5, each normalizing x to [0, 1]. Model selection used the Akaike Information Criterion (AIC). Exons with a negative linear slope (decreasing NMD efficiency towards the 3′ end of the exon) were excluded from PCA.

### Principal component analysis

#### For start-proximal DMS

PCA was applied to the gene × position LOESS-interpolated matrix (83 positions, see above). Each position (feature) was standardized by z-score normalization (zero mean, unit variance; ‘StandardScaler’) across genes before PCA. The resulting principal components capture gene-level variation in the shape of the NMD efficiency profile across the first 83 PTC positions.

#### For long exon data

PCA was applied to interpolated NMD efficiency curves from long exons (>400 nt, excluding first, last, and penultimate coding exons). For each exon, predictions were linearly interpolated at 50 uniformly spaced relative positions (0–1 within the exon). Exons with fewer than 5 data points or with invalid (NaN/Inf) interpolated values were excluded. Features were z-score normalized before PCA. Three principal components were retained.

#### Hierarchical clustering of start-proximal genes

Ward-linkage hierarchical clustering was applied to the sigmoid parameters (A, K, B, M, R²) derived from per-gene 4-parameter sigmoid fits (see above). Outlier genes (*FAT1*) were removed prior to clustering. Parameters were z-score standardized before computing the Euclidean distance matrix. The optimal number of clusters was evaluated by silhouette score for k = 2–7; k = 3 was selected. The same clustering approach was applied to genome-wide start-proximal predictions, using sigmoid parameters fitted to NMDetective-AI predictions at positions 1–83 codons. Genes were retained for genome-wide clustering only if they had at least 10 data points in the start-proximal window, a first CDS exon ≤ 400 nt, a CDS ≥ 500 nt, and the start-proximal window did not reach within 55 nt of the last exon junction. Cluster enrichments for gene categories (essential genes, tumor suppressors, oncogenes, disease genes) were assessed by Fisher’s exact test.

#### SpliceAI scores

We used SpliceAI to predict splicing site donors and acceptors along the full transcripts (2958 nts), comprising the variable (library) and constant (*BRCA1*) parts. SpliceAI outputs a donor and acceptor likelihood for each position of the transcript. It correctly predicted the three splice events in the constant part of the transcript, validating the analysis. For a minority of genes, it also predicted some splice sites within the library. To check whether these genes were enriched in the NMD-resistant group of genes we computed the SpliceAI score, which results from summing the splice site likelihood over all library nucleotides. This metric represents a likelihood of observing alternative splice events involving library nucleotides.

### Selection in germline and soma

#### Germline variant data and NMD classification

Stop-gained variants from gnomAD v4.1 (https://gnomad.broadinstitute.org/news/2024-04-gnomad-v4-1/) were annotated with NMD status categories. Rare germline variants were defined as those with allele frequency (AF) < 0.1% across all gnomAD subpopulations (African, Latino/Admixed American, Finnish, Non-Finnish European, East Asian), while common variants had AF ≥ 0.1% in at least one subpopulation. Only PASS-filtered variants with VEP stop_gained annotations in MANE Select transcripts were retained. Variants were classified as NMD-evading if they met any of four criteria: within the first 150 coding nucleotides from the start codon (start-proximal), within 55 nucleotides upstream of the last exon-exon junction (55-nucleotide rule), within the last coding exon, or within an exon longer than 400 nucleotides (long exon). Variants meeting none of these criteria were classified as NMD-triggering.

#### Optimizing NMDetective-AI prediction thresholds

For NMDetective-AI-based classification, NMDetective-AI predictions were generated for all variants using full-length transcript sequences encoded as 6-track representations (see above) obtained via GenomeKit (GENCODE v47). To establish thresholds for binarizing predictions, a 3-component Gaussian mixture model (GMM) was fitted to prediction distributions from rare gnomAD variants using scikit-learn’s GaussianMixture with full covariance. The three components correspond to NMD-evading, intermediate, and NMD-triggering categories. Thresholds were defined as the intersection points between adjacent Gaussian components, computed by finding the roots of the difference between weighted probability density functions using Brent’s method. This yielded thresholds of −0.17 (evading vs intermediate) and 0.43 (intermediate vs triggering). Variants with predictions in the intermediate range were excluded from binary classification analyses.

#### Per-gene NMD variant composition in germline

Gene-level log odds ratios were calculated as log_e_[(N_triggering + 1) / (N_evading + 1)], where pseudocounts avoided division by zero. Genetic constraint was assessed using LOEUF (loss-of-function observed/expected upper bound fraction) deciles from gnomAD, with highly constrained genes defined as LOEUF decile 0–1.

To assess NMD variant enrichment in ClinVar disease genes, two complementary Wald log OR tests were computed per gene (with Haldane-Anscombe 0.5 pseudocounts). Test 1 (T1) compares rare PTC allele counts in NMD-triggering versus NMD-evading regions, normalized by synonymous (plus missense) allele counts in the same regions, restricted to PTC variants with AF ≤ 1% to avoid extreme-AF variants dominating. Test 2 (T2) provides an internal control: within each gene, PTC variants are ranked by AF and split at the gene-level median into a rarer half and a more common half; the 2×2 table compares NMD-triggering versus NMD-evading counts in rare versus common within-gene halves. A combined *p*-value was derived via Fisher’s method (−2·Σ log(p) ∼ χ² (4)). P-values for each test were corrected independently using Benjamini-Hochberg FDR. Only disease genes with ≥10 unique PTC variants and non-zero synonymous counts in each NMD region were included. Genes were considered robustly enriched when passing T1 FDR < 1% and T2 FDR < 25%.

#### Somatic PTC analysis: gene-category meta-analysis

To investigate selection patterns in somatic mutations, we analyzed somatic PTCs from TCGA ∼17k whole-genome sequences (v43) alongside synonymous variants from the same samples as a mutation-rate control. Variants were classified into NMD-triggering and NMD-evading regions using the same rule-based criteria as for germline variants. Synonymous variants were assigned NMD status based on their genomic position relative to the same exon boundaries (i.e., whether a hypothetical PTC at that position would be NMD-triggering or NMD-evading), providing a positional control for regional mutation rate variation. For each gene, a 2×2 contingency table was constructed:

|  | NMD-triggering | NMD-evading |
| --- | --- | --- |
| PTCs | a | b |
| Synonymous | c | d |

A Wald-adjusted log odds ratio was computed as loge OR = log_e_[(a + 0.5)(d + 0.5) / (b + 0.5)(c + 0.5)], where a pseudocount of 0.5 was added to all cells. The standard error was estimated as SE = √(1/(a+0.5) + 1/(b+0.5) + 1/(c+0.5) + 1/(d+0.5)), and a two-sided z-test yielded the per-gene P-value. Positive log OR indicates that PTCs in that gene are disproportionately concentrated in NMD-triggering positions relative to what synonymous mutations would predict, consistent with positive selection for NMD-mediated loss of function.

Genes were assigned to four categories: tumor suppressor genes (TSGs) and oncogenes (OGs) from curated lists of mutational cancer drivers^68,69^, essential genes from^70^, and other genes. To ensure a clean background set for comparisons, “Other Genes” excluded all genes appearing in any row of the cancer gene census (not only common drivers) and all genes with disease associations in the NCBI gene_condition_source_id file (https://ftp.ncbi.nlm.nih.gov/pub/clinvar/gene_condition_source_id). Only genes with ≥10 PTCs and ≥5 synonymous variants were included.

Per-gene log ORs within each category were pooled using inverse-variance–weighted fixed-effects meta-analysis. Heterogeneity was assessed by Cochran’s Q statistic and I². When substantial heterogeneity was detected (I² > 50%), a DerSimonian-Laird random-effects model was used instead. Each category was compared against the “Other Genes” background via a z-test on the difference of pooled log ORs, with SE computed as √(SE²_category + SE²_other).

As a sensitivity analysis, the classification was repeated with an NMDetective-AI filter. For each gene with ≥10 PTCs with NMDetective-AI predictions, a two-component GMM was fitted to the predictions to find a gene-specific decision boundary (the prediction value where both components had equal posterior probability). Variants where NMDetective-AI prediction disagreed with the rule-based classification (NMDetective-AI says triggering but rules say evading, or vice versa) were marked as unknown and excluded. This filter removes ambiguous variants to test whether the selection signal is robust to classification uncertainty.

#### Per-gene analysis of individual cancer genes

For individual cancer gene analysis, the same Wald log OR approach was applied to each tumor suppressor gene (including both common and rare mutational drivers). Genes with ≥3 unique PTCs and ≥3 synonymous variants in each NMD region were retained. *P*-values were corrected for multiple testing using Benjamini-Hochberg false discovery rate (FDR). Genes with FDR < 0.05 were considered significant.

## Code and data availability

All the code related to training NMDetective-AI, dataset processing, downstream analysis, and generating figures of this article can be found at https://github.com/Vejni/NMDetectiveAI. Model weights, analyzed datasets and source data files for all figures can also be downloaded from the same repository. Filtered and processed PTC variants and NMD efficiency estimates are available upon request. Precomputed NMDetective-AI predictions for all stop-gain mutations along the longest transcripts of each gene can be found on the above repository.

## Acknowledgements

M.V. was supported by a fellowship from the ”la Caixa” Foundation (ID 100010434, with fellowship code B006197). G.P.-M. was funded by an AGAUR FI fellowship.

Work in the lab of F.S. is supported by an ERC StG “HYPER-INSIGHT” (757700), an ERC Consolidator Grant “STRUCTOMATIC” (101088342), Horizon2020 RIA project “DECIDER” (965193), Horizon Europe project “LUCIA” (101096473), Spanish government project “REPAIRSCAPE”, CaixaResearch project “POTENT-IMMUNO” (HR22-00402), a Novo Nordisk Fonden “Start Package” grant, the Danish Cancer Society grant “AI-DRIVERS” and a DFF Project2 (5243- 00072B), the SGR funding of the Catalan government, and the Severo Ochoa Centers of Excellence award of the Spanish government to the IRB Barcelona.

Work in the lab of B.L. was funded by European Research Council (ERC) Advanced (grant 883742), Wellcome (220540/Z/20/A), the Spanish Ministry of Science and Innovation (PID2023-146685NB-I00, EMBL Partnership, Severo Ochoa Center of Excellence), Agencia de Gestio d’Ajuts Universitaris i de Recerca (AGAUR, 2021-SGR-01226) and the CERCA Program/Generalitat de Catalunya.

LLM-assisted tools were used for language editing of manuscript text. The results published here are in whole or part based on data generated by the TCGA Research Network (https://www.cancer.gov/tcga).

