## Supplementary Figures S1-S17 for "Quantitative prediction of nonsense-mediated mRNA decay across human genes by genomic language model and large-scale mutational scanning"


- Figure S1: Filtering steps on TCGA and GTEx nonsense variants.
- Figure S2: Additional filtering and preprocessing steps for modeling.
- Figure S3: Distribution of processed NMD efficiency scores.
- Figure S4: Hyperparameter optimization via Bayesian hyperparameter search.
- Figure S5: Interreplicate correlation for DMS experiments.
- Figure S6: RNA level observations from the penultimate exon DMS experiment.
- Figure S7: Evasion patterns of penultimate exon PTCs.
- Figure S8: RNA level observations from the long exon DMS experiment.
- Figure S9: RNA levels across exon lengths and proposed model.
- Figure S10: NMD efficiency in long exons across DMS experiment, genomic data and predictions.
- Figure S11: Predicted NMD efficiency profiles of example long exons from the genome.
- Figure S12: NMD efficiency scores of the start proximal DMS experiment before scaling.
- Figure S13: NMD efficiency scores of the start proximal DMS experiment after rescaling.
- Figure S14: Start proximal NMD evasion shape clusters in the genome.
- Figure S15: Impact of PTC Position and Stop Codon Type on RNA Levels.
- Figure S16: Finding the optimal NMDetective-AI prediction threshold from population variants.
- Figure S17: Correlation between selection tests of NMD triggering vs evading PTC bearing disease genes.


**Supplementary Tables:**


- Table S1: Penultimate exon 4PL parameters and performance for all genes.
- Table S2: PC values for all analyzed long exons.
- Table S3: Start proximal sigmoid fits and cluster assignments for all genes.
- Table S4: Selection test results for tested disease genes.

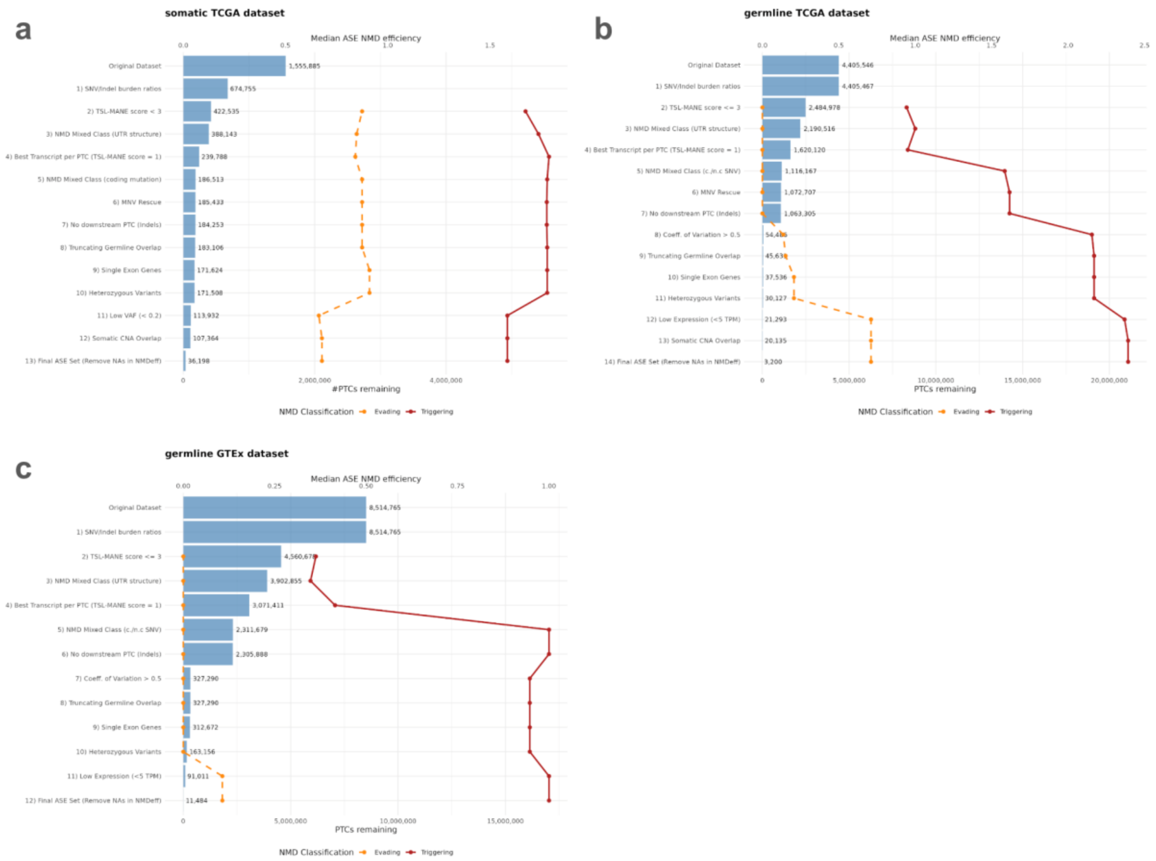


**Figure S1: Filtering steps on TCGA and GTEx nonsense variants.**

The y-axis shows the different filtering steps performed on somatic (a) and germline (b) TCGA PTC variants, and GTEx (c). The bars show the number of variants at each point in the preprocessing, lineplots denote the means of ASE based NMD efficiency estimates of evading (yellow) and triggering (red) variants based on known rules of NMD.

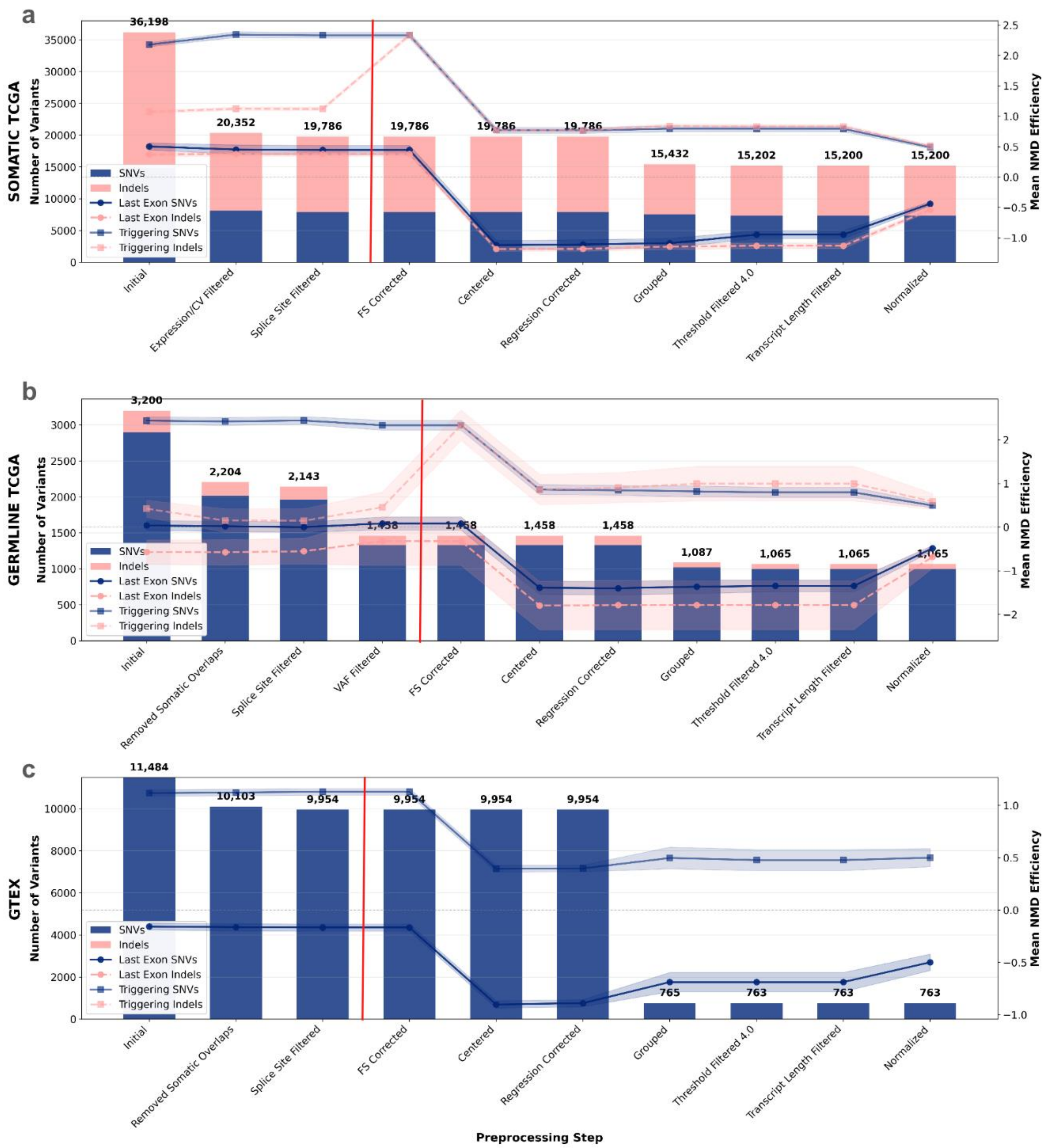


**Figure S2: Additional filtering and preprocessing steps for modeling.**

The above plots show additional preprocessing steps on the PTC variant sets of somatic TCGA (a), germline TCGA (b), and GTEx (c). Heights of bars show the number of variants at that point of the preprocessing with insertion-deletion variants in pink and SNVs in blue. Lineplots in matching colours also denote the means of the respective variant types that trigger NMD (square), and evade NMD (circle, only last exon corresponding to full evasion). The read vertical lines on each panel denote the point where Files 1-3 were saved at, while Files 4-6 were saved at the end of the processing pipeline. For explanation of steps see Methods.

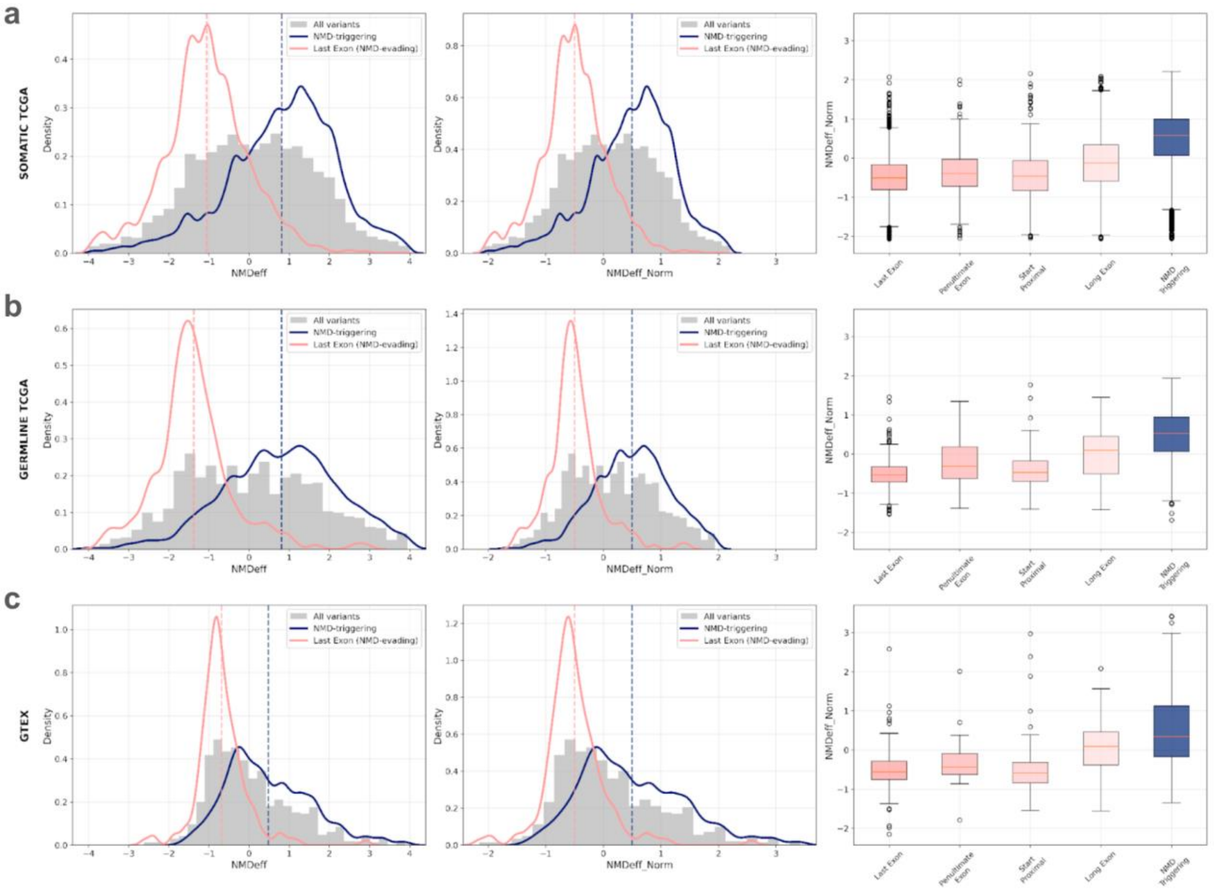


**Figure S3: Distribution of processed NMD efficiency scores.**

Plots show the distribution of processed NMD efficiency (NMDeff) scores across datasets: somatic TCGA variants (a), germline TCGA variants (b), GTEx (c). First column, NMDeff, denotes the variantsets before the last point in the previous figure, while the second column (NMDeff_Norm) denotes variants normalized to the scale of -0.5 (full evasion) to 0.5 (full triggering). KDEs show the distribution of NMD triggering (blue), and last exonic (full evading) variants. Boxplots in the third column show the NMDeff_Norm distribution of variants classified according to different NMD rules.

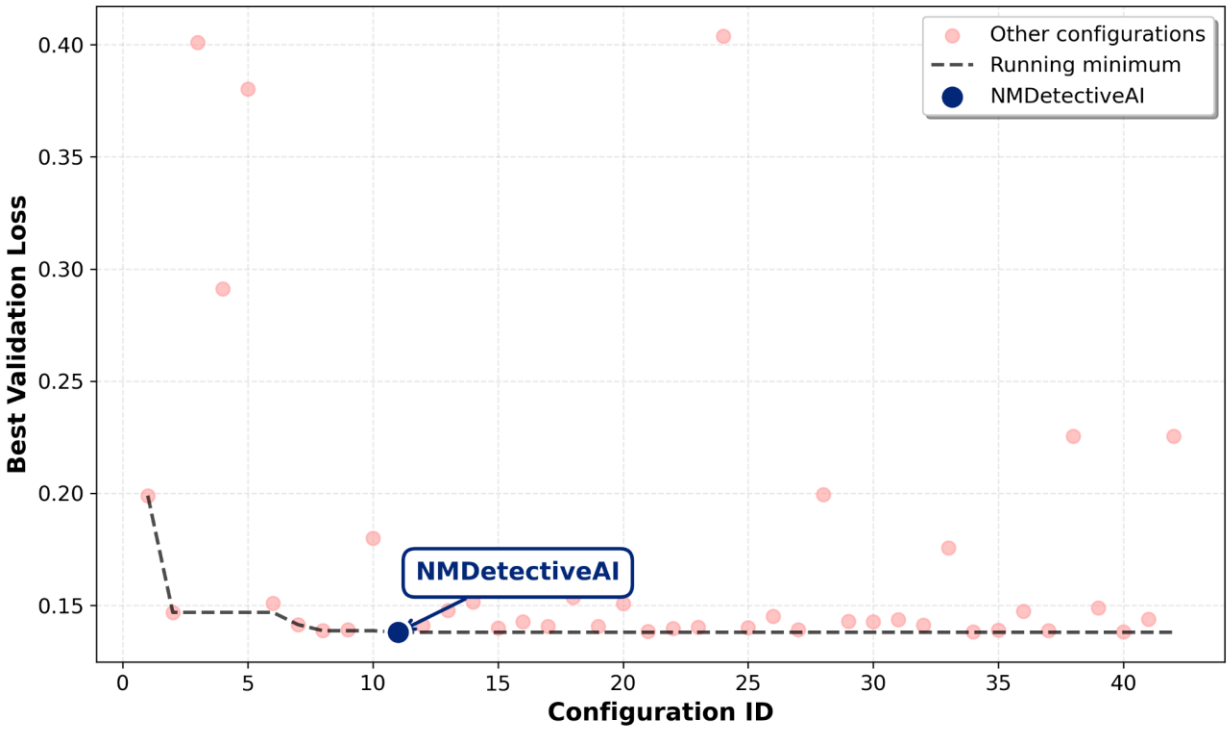


**Figure S4: Hyperparameter optimization via Bayesian hyperparameter search.**
Model performance, measured as the lowest validation set loss (y-axis), across different hyperparameter sets (x-axis). The model with best performance across the search was kept ad NMDetective-AI, and is annotated on the plot.

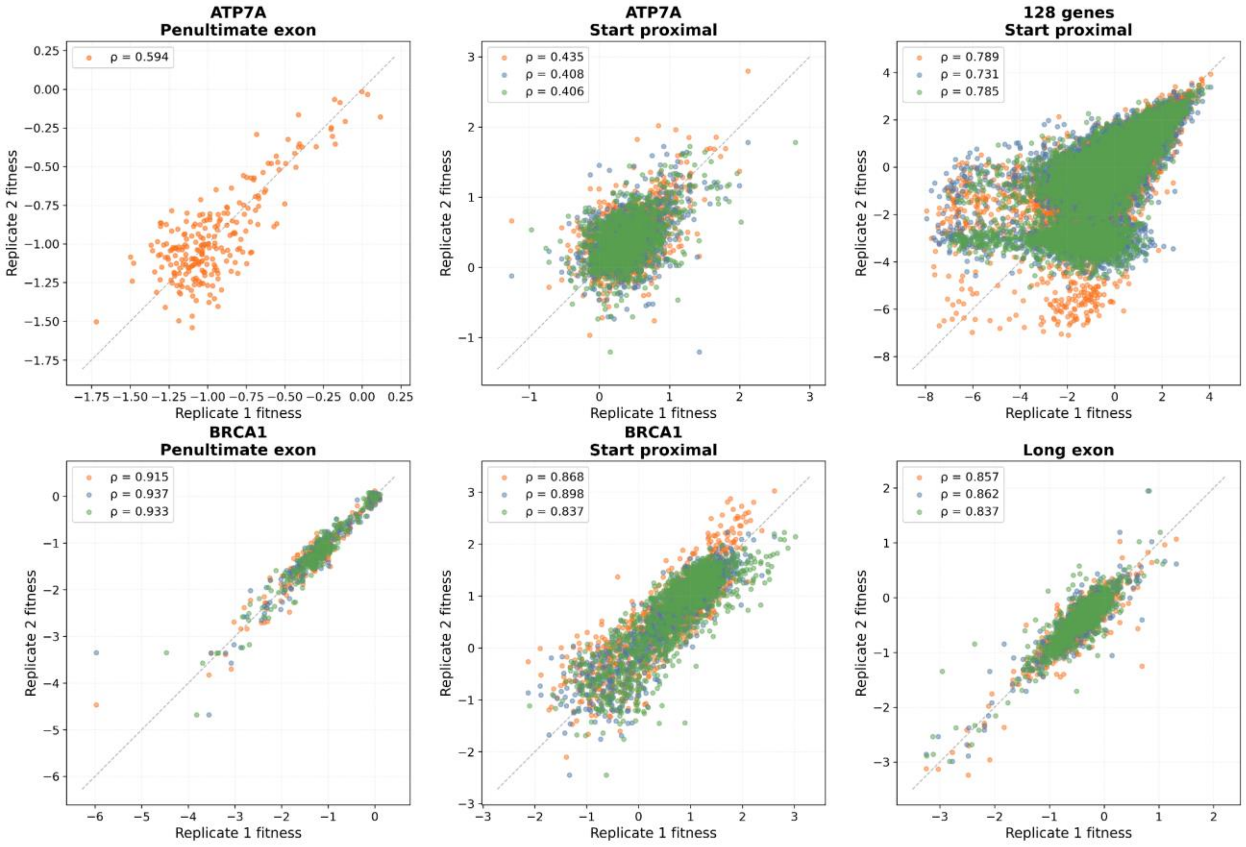


**Figure S5: Interreplicate correlation for DMS experiments.**

Grid of scatter plots showing biological replicate comparisons for 6 DMS experiments (50 nt/penultimate exon experiment in *ATP7A, and BRCA1* minigene (first column). Start proximal experiments in *ATP7A, and BRCA1 (second column).* Start proximal experiment across 139 genes (122 high quality genes shown; third column, top row). Long exon experiment in *BRCA1* (third column, bottom row). All three pairwise comparisons of 3 biological repeats are overlaid in different colors (orange: rep1 vs rep2; blue: rep1 vs rep3; green: rep2 vs rep3). Spearman correlation coefficients (ρ) shown in legends. Dashed black line indicates perfect correlation.

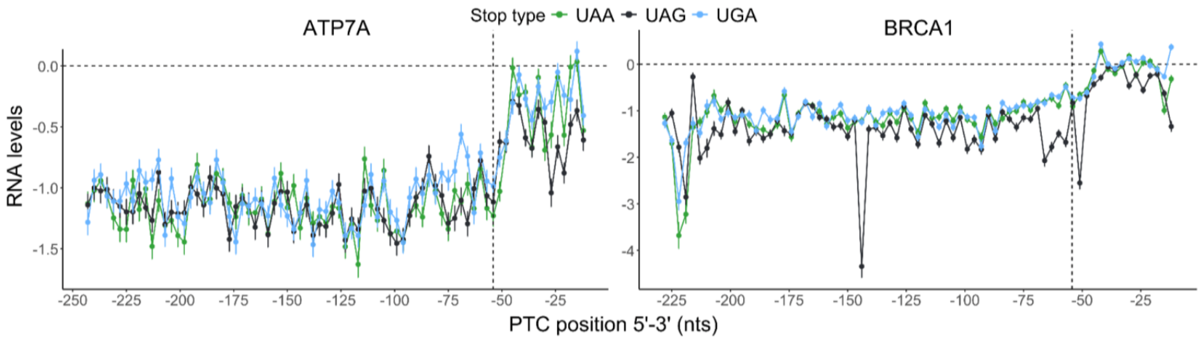


**Figure S6: RNA level observations from the penultimate exon DMS experiment.**
RNA levels for ATP7A (left) and BRCA1 (right) PTC variants along the 5'-3' axis and coloured by stop type. The x-axis is in nucleotide units, and -1 is the first position upstream of the EJC. Scores of 0, <0 and >0 indicate equal, lower and higher RNA levels than WT; respectively. Error bars indicate the DiMSum-calculated error across replicates. The vertical dashed line indicates the residue spanning nucleotide positions 54-57 upstream of the EJC, which represents the boundary of the 50-55 nts rule as described in literature. The horizontal dashed line represents the wild-type RNA levels.

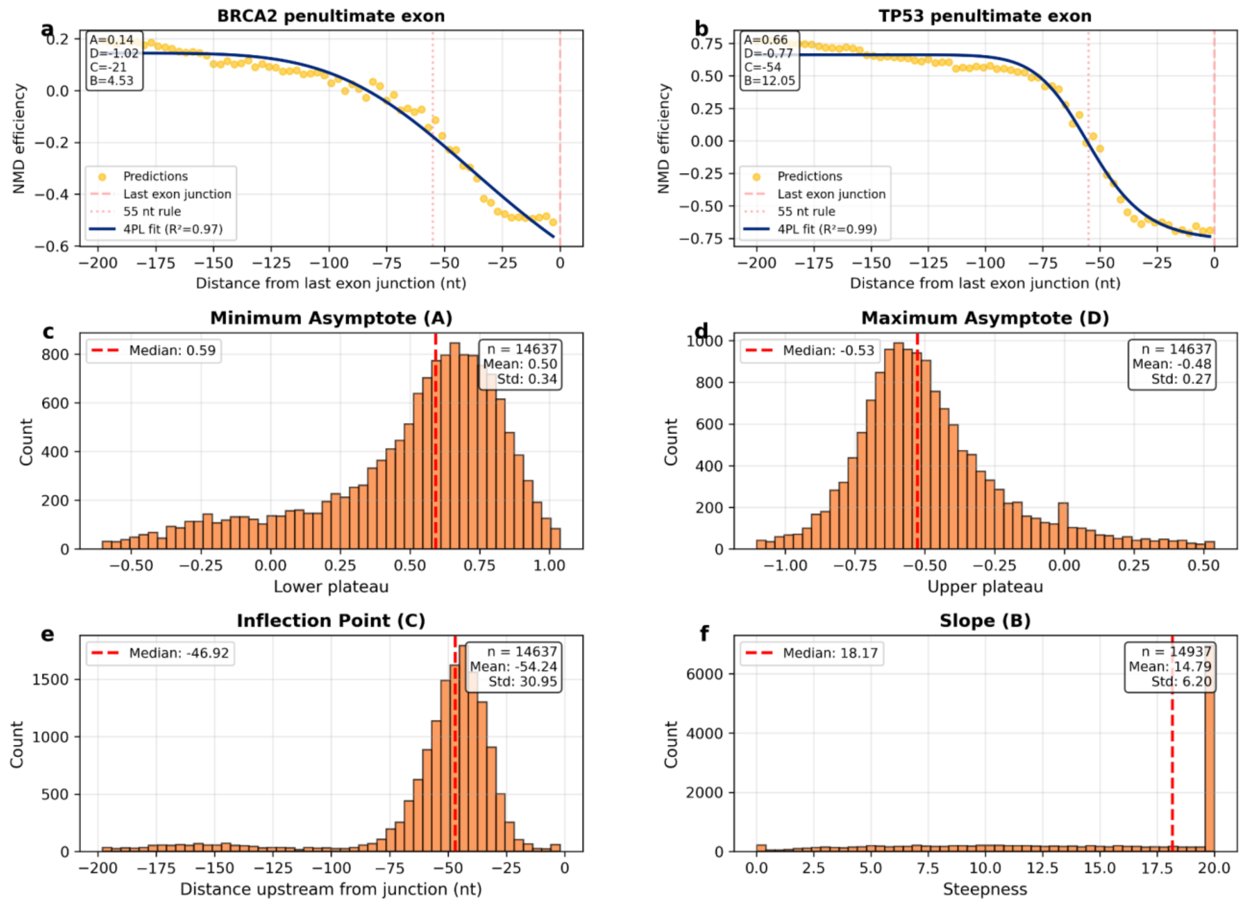


**Figure S7: Evasion patterns of penultimate exon PTCs.**

Top row denotes NMDetective-AI prediction NMD efficiency scores of PTCs inserted at penultimate exon positions of the *BRCA2* and *TP53* genes. Vertical lines denote the last exon boundary and 55nt upstream of the last exon, denoting the 55nt NMD evasion rule. 4PL fits to predictions shown in blue, with the parameters denoted on the plot (top left corner). Histograms of the distributions of 4PL parameters fit to all coding genes genome-wide.

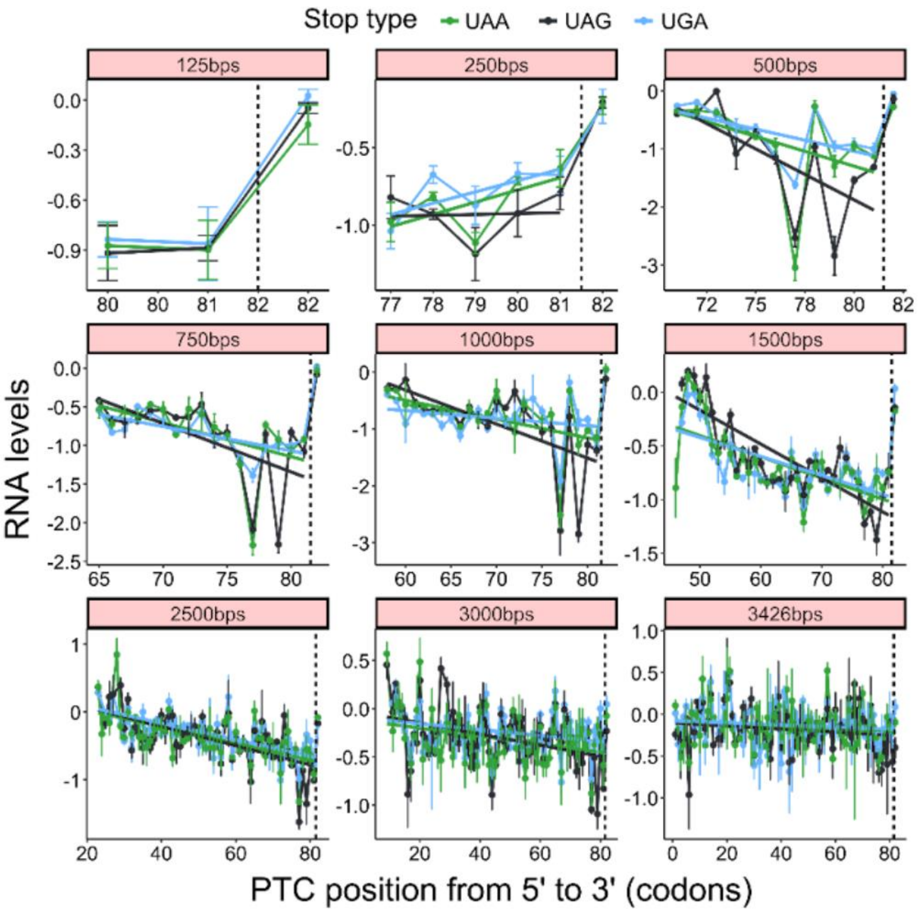


**Figure S8: RNA level observations from the long exon DMS experiment.**

RNA levels for the PTC variants along the 5'-3' axis for each exon length (quadrant) and coloured by stop type. Scores of 0, <0 and >0 indicate equal, lower and higher RNA levels than the UGG WT; respectively. Color lines show the correlation between PTC position and RNA levels for each stop type, excluding PTC 82 because it is already rendered NMD resistant by the 50-55 nts rule. The vertical dashed line indicates the residue spanning nucleotide positions 54-57 upstream of the EJC, which represents the boundary of the 50-55 nts rule. Error bars indicate the DiMSum-calculated error across replicates.

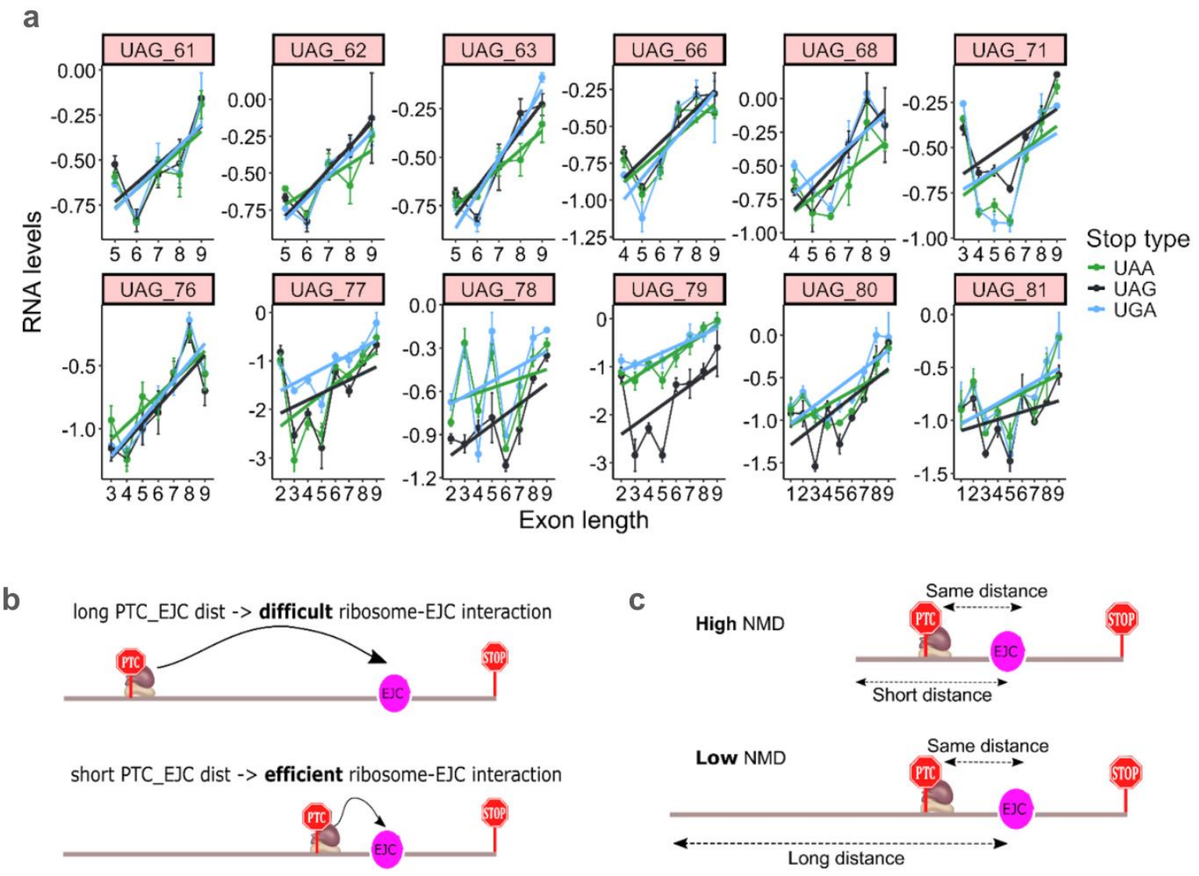


**Figure S9: RNA levels across exon lengths and proposed model.**

All data points in each plot of on panel (a) have the same PTC-to-downstream intron distance, but different sequence lengths upstream of the PTC. X-axis numbers represent exon lengths as indicated in a. Scores of 0, <0 and >0 indicate equal, lower and higher RNA levels than WT; respectively. Color lines show the correlation between PTC position and RNA levels for each stop type. Color lines show the correlation between the exon length and the RNA levels for each stop type. Error bars indicate the DiMSum-calculated error across replicates. (b) Representation of the PTC-to-downstream intron distance NMD model, where increased distances hinder the recruitment and assembly of the NMD machinery due to weaker interactions between the ribosome-bound UPF1 and the EJC. (c) Representation of the exon length effect on NMD.

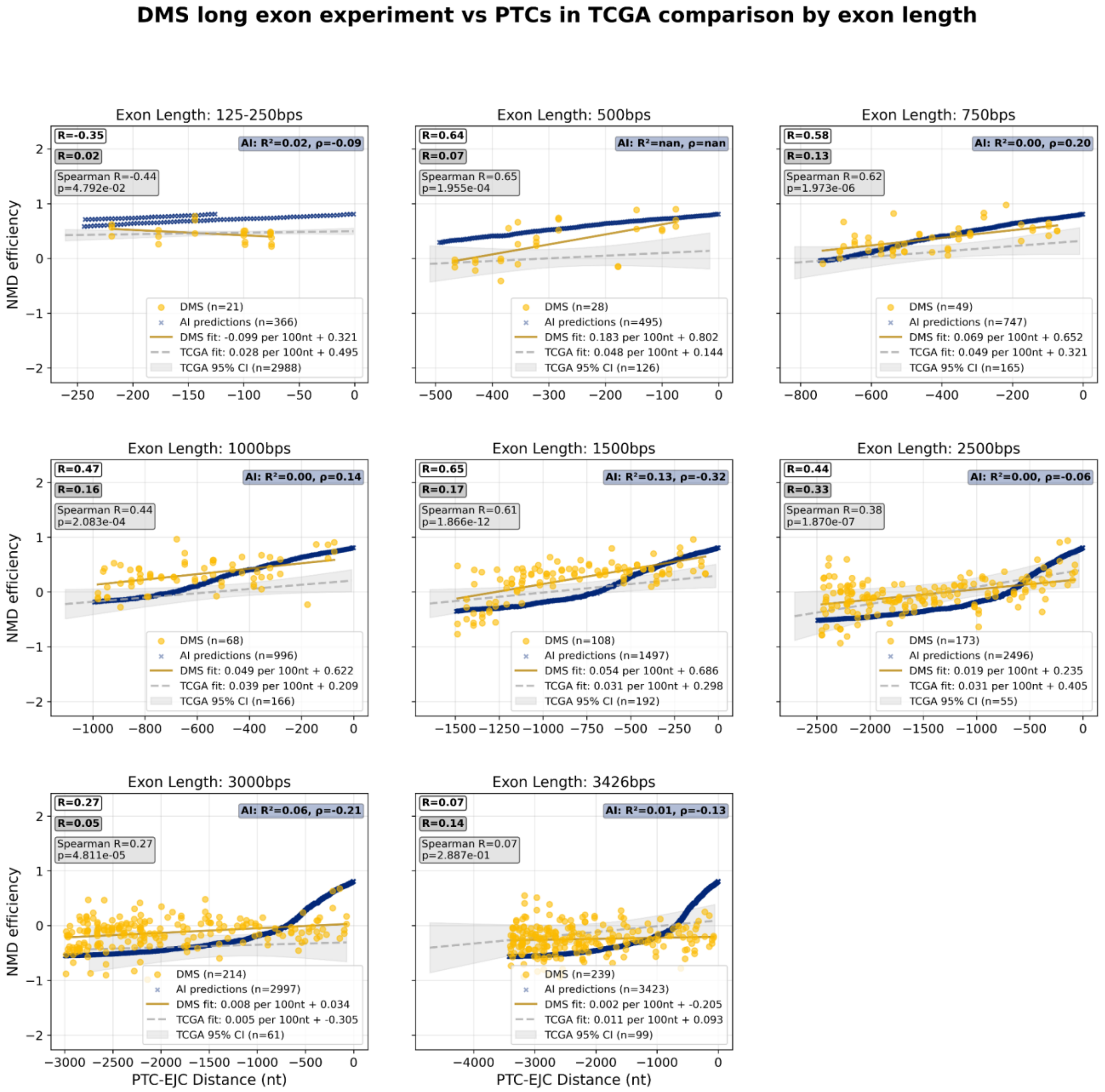


**Figure S10: NMD efficiency in long exons across DMS experiment, genomic data and predictions.**

Comparison of NMD efficiency between DMS long exon experiment (yellow circles) and TCGA PTCs (gray squares) across all exon length categories. Each panel shows scatter plots with linear regression fits (solid lines) and correlation coefficients. NMDetective-AI predictions shown as crosses. The x-axis shows distance from the exon-exon junction (negative values indicate upstream positions).

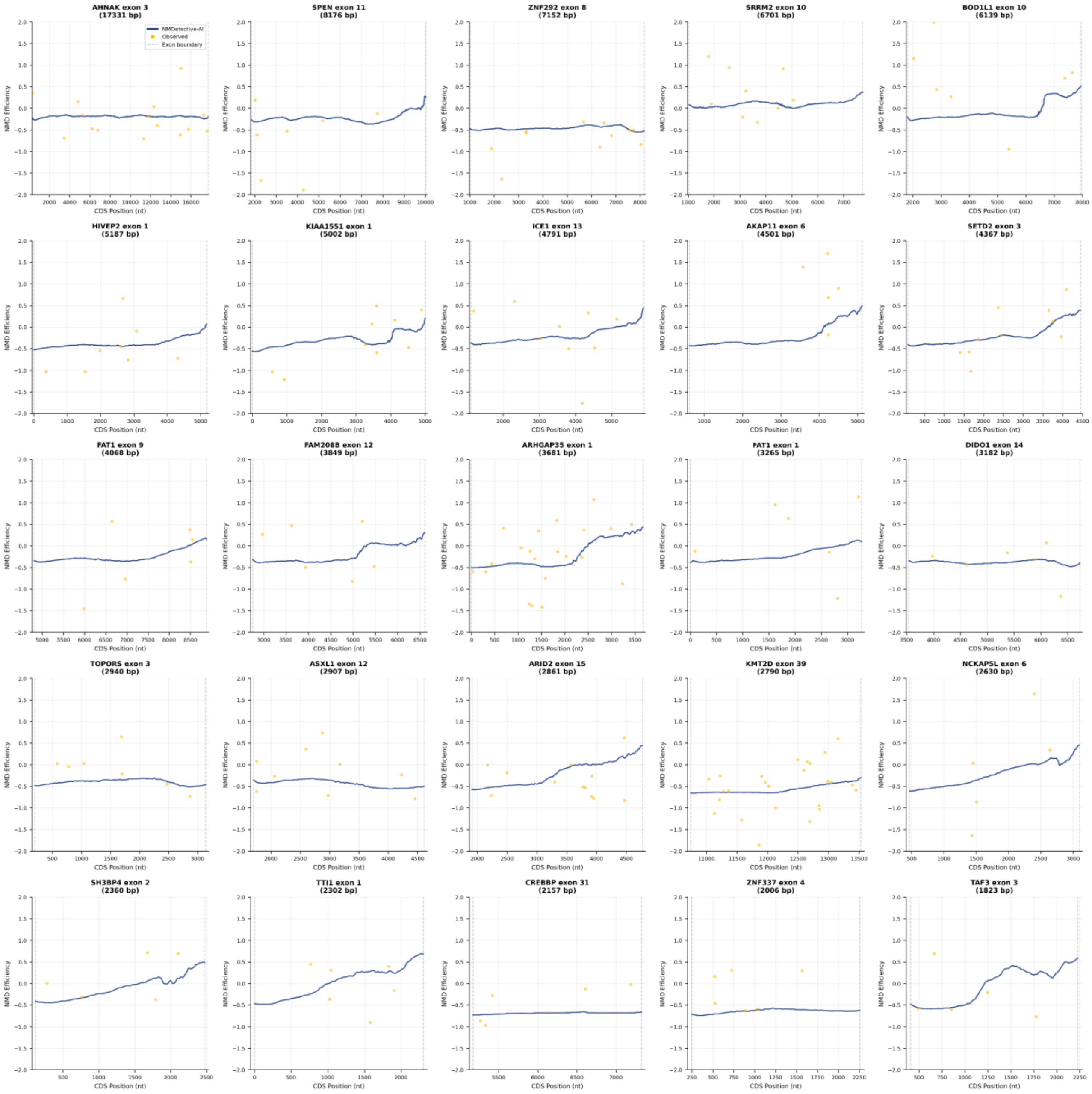


**Figure S11: Predicted NMD efficiency profiles of example long exons from the genome.** NMDetective-AI predictions (blue, y-axis) of PTCs inserted in various positions (x-axis) of long exons in the genome. Gene name, exon number, as well as exon length are shown on the top of each plot. The longest transcript of each gene was considered. Observations of somatic PTCs in TCGA are shown as yellow dots.

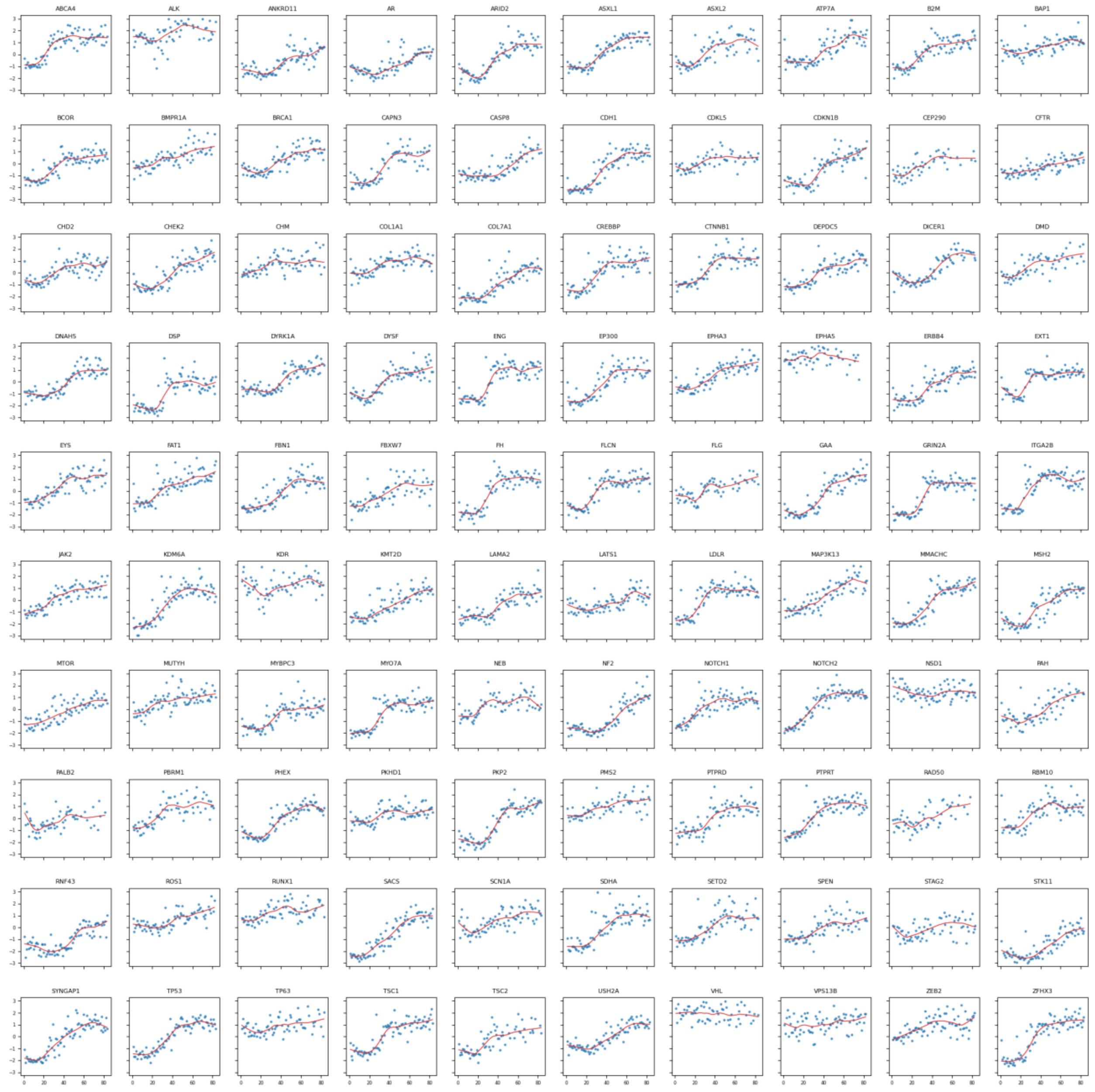


**Figure S12: NMD efficiency scores of the start proximal DMS experiment before scaling.** RNA level observations from the start proximal DMS experiment were sign-flipped and centered around 0 by subtracting the mean across the whole dataset. Shown for high quality (n=100) genes. Observations shown as blue dots (y-axis), while the red line denotes LOESS fits per gene, across PTC positions (x-axis).

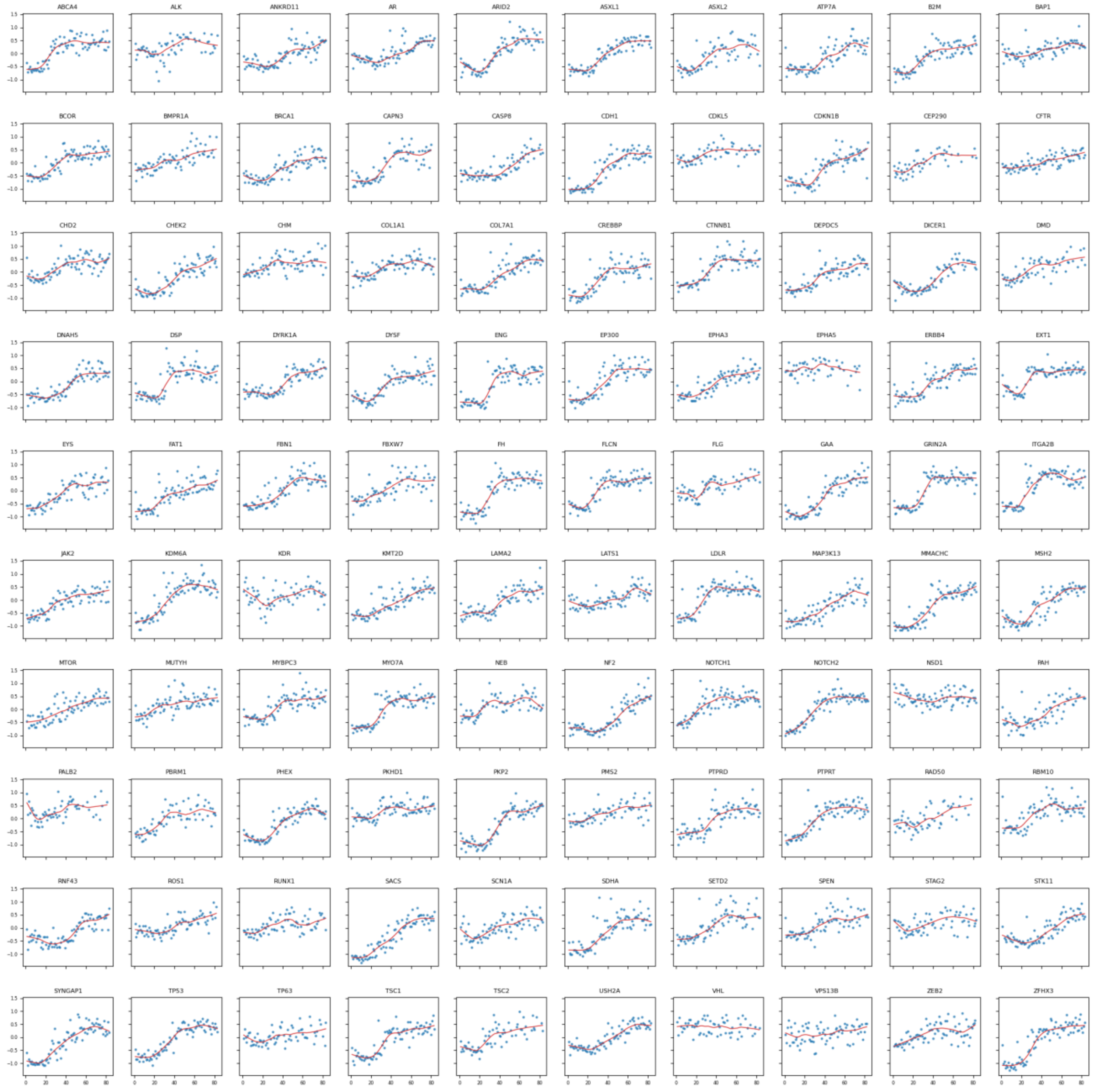


**Figure S13: NMD efficiency scores of the start proximal DMS experiment after rescaling.** NMD efficiency scores from the start proximal DMS experiment were rescaled to the proposed NMDeff scale (-0.5 full evasion, 0.5 full NMD triggering), while maintaining inter-gene variability at the 5' end of genes (see Methods). Shown for high quality (n=100) genes. Observations shown as blue dots (y-axis), while the red line denotes LOESS fits per gene, across PTC positions (x-axis).

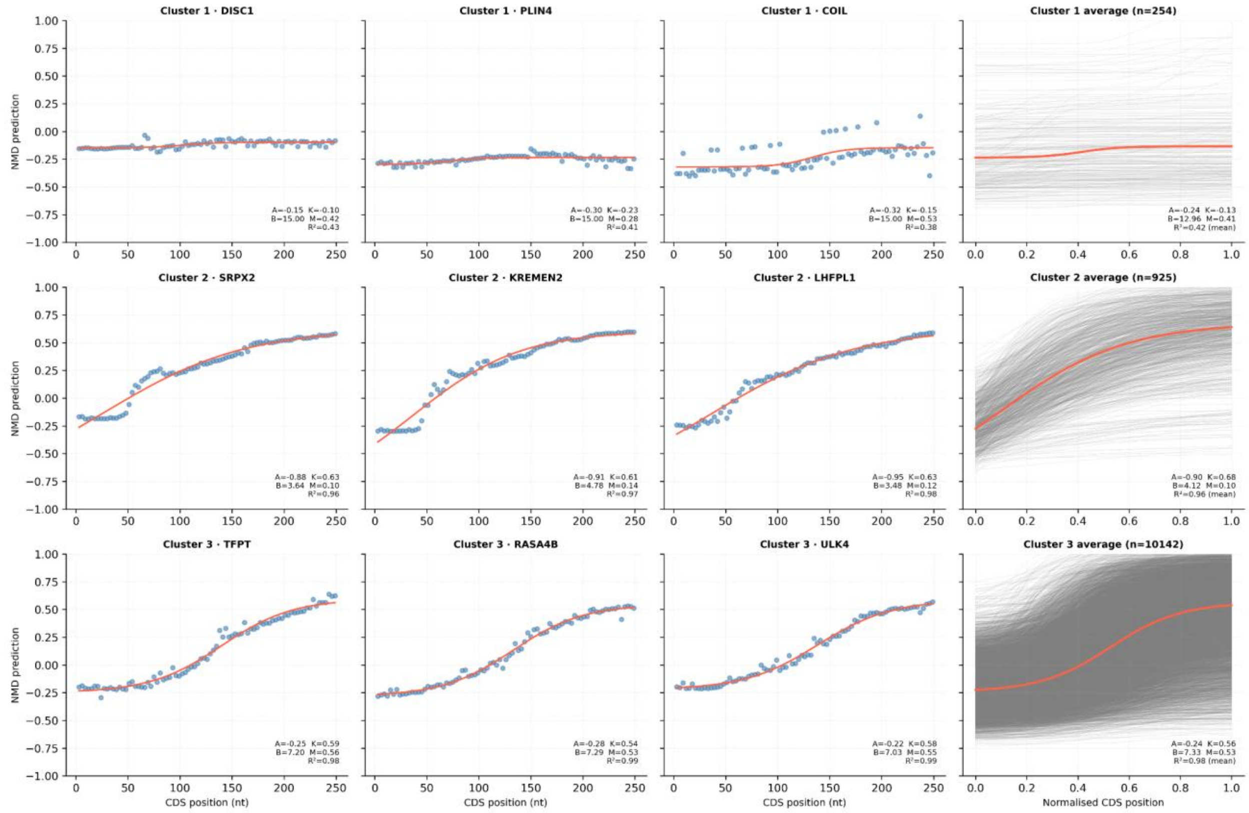


**Figure S14: Start proximal NMD evasion shape clusters in the genome.**
4 parameter sigmoid curves were fit to PTC scan predictions from NMDetective-AI genome-wide and clustered using hierarchical clustering. A clustering with 3 clusters were considered. 3 example start proximal regions are shown per cluster. NMDetective-AI predictions are shown as blue dots, with the 4 parameter sigmoid fit in red. The distribution and average sigmoid curves of that cluster are shown in the fourth column.

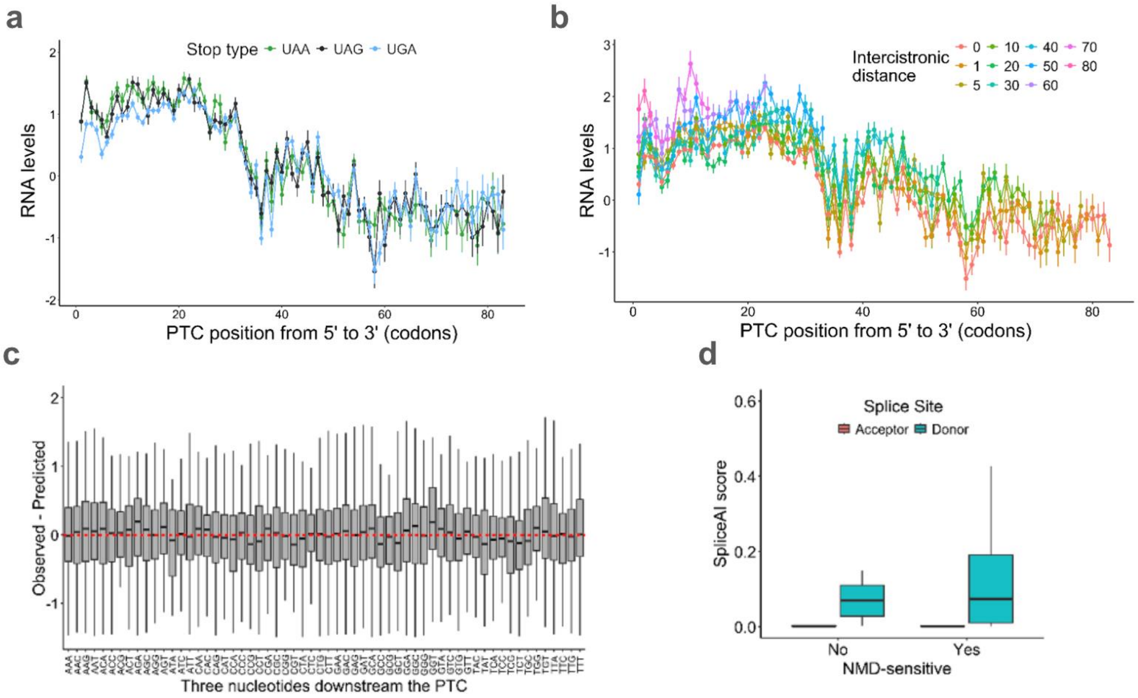


**Figure S15: Impact of PTC Position and Stop Codon Type on RNA Levels.**
(a) RNA levels for non-AUG variants as a function of PTC position along the 5'-3' axis and coloured by stop type. Scores of 0, <0 and >0 indicate equal, lower and higher RNA levels than WT; respectively. Error bars indicate the DiMSum-calculated error across replicates. (b) Same as in (a) but now also including the variants with AUGs downstream of the PTC. Only UGA variants are shown. The color legend shows the PTC-AUG distance (intercistronic distance). (c) Histograms showing the error in the RNA levels predicted by the loess start proximal model for different sequence features, being codon downstream of the PTC codon. (d) For each gene, we calculated the SpliceAI score by summing the splice site likelihood over all the nucleotides of the library (SpliceAI score), and correlated it with the absence and presence of NMD-sensitivity in genes. NMD-insensitive genes are those where all PTC positions have equal or higher RNA levels than WT (shown in a, top).

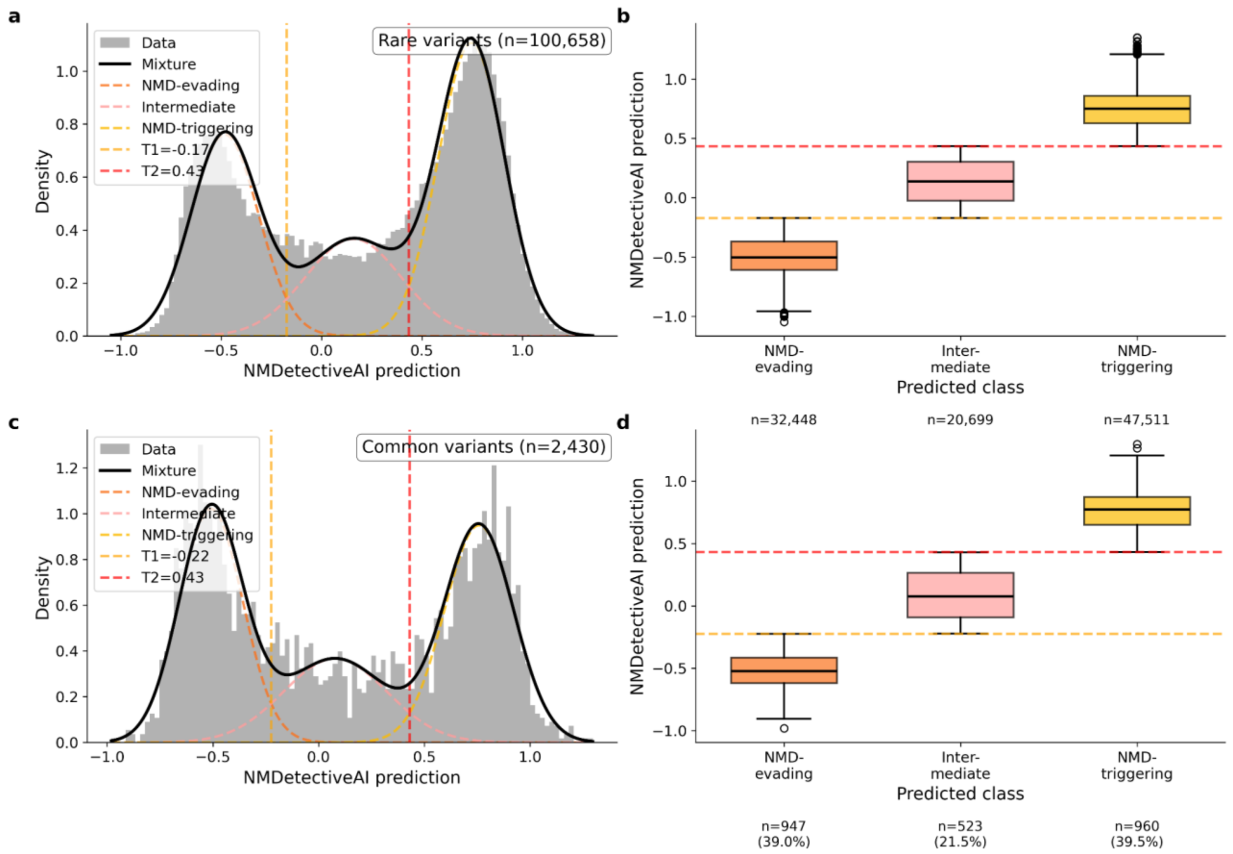


**Figure S16: Finding the optimal NMDetective-AI prediction threshold from population variants.**

Distribution of NMDetective-AI predictions of rare (a) and common (b) PTCs in GnomAD. The three distributions fit by a Gaussian Mixture Model, and the two thresholds best separating them are shown as dashed lines. Boxplots in the second column show the proportion of variants in each group: predicted NMD evading, intermediate and NMD triggering.

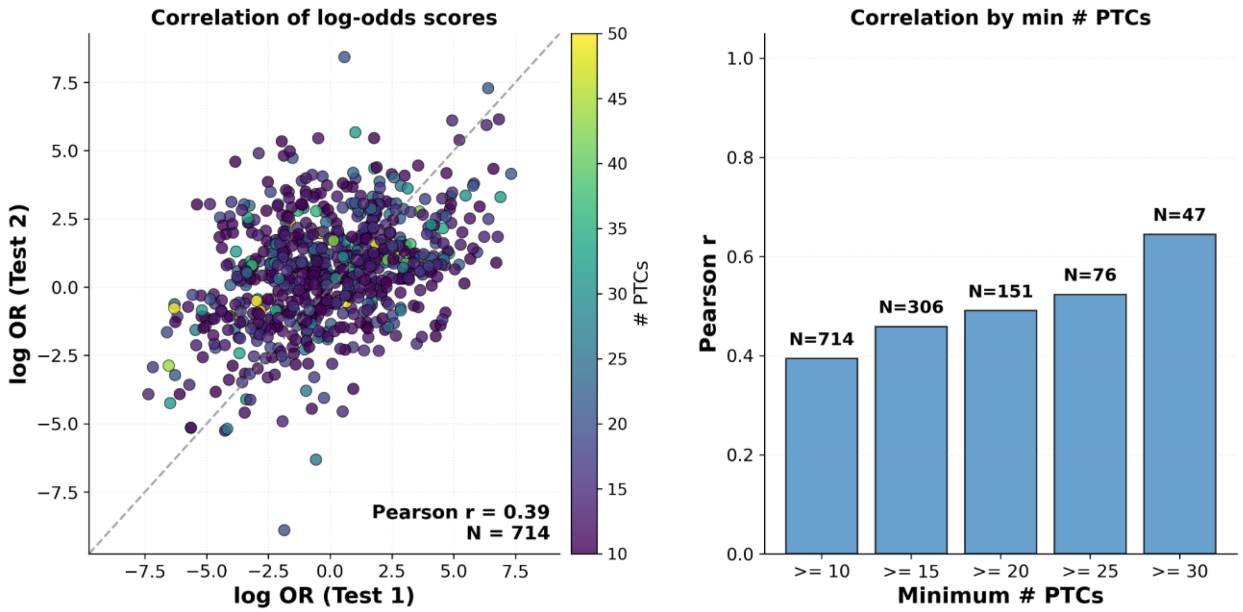


**Figure S17: Correlation between selection tests of NMD triggering vs evading PTC bearing disease genes.**

Log odds ratio values of NMD triggering vs NMD evading PTCs, normalized by the synonymous and missense mutation counts in the same genes, calculated from two different analyses. Each dot represents a gene, with >= 10 PTC variants (see Methods for other filters). Test 1 (T1) compares rare PTC allele counts in NMD-triggering versus NMD-evading regions, normalised by synonymous allele counts in the same regions; Test 2 (T2) uses a within-gene internal control, comparing rarer versus more common PTCs within each gene. The barplot on the right shows the Pearson correlation between Test 1 and Test 2 increasing as genes are filtered more heavily by number of PTCs.
